# Bacterial microbiota of *Ostreobium*, the coral-isolated chlorophyte ectosymbiont, at contrasted salinities

**DOI:** 10.1101/2023.01.31.526156

**Authors:** A. Massé, J. Detang, C. Duval, S. Duperron, A.C. Woo, I. Domart-Coulon

**Affiliations:** Molécules de Communication et Adaptation des Microorganismes (MCAM), Muséum national d’Histoire naturelle (MNHN), CNRS (UMR7245); CP54, 63 Rue Buffon, 75005, Paris, France; Pôle Analyse de Données UAR 2700 2AD, Muséum national d’Histoire naturelle (MNHN); 43 Rue Cuvier, 75005, Paris, France

**Keywords:** Ulvophyceae, bacterial communities, microbiota, salinity, 16S rDNA metabarcoding, CARD-FISH

## Abstract

Microscopic filaments of the siphonous green algae *Ostreobium* (Ulvophyceae, Bryopsidales) colonize and dissolve the calcium carbonate skeletons of coral colonies, in shallow-water reef environments of contrasted salinities. Their bacterial composition and plasticity in response to salinity remain unknown. Here, we analyzed the bacteria associated with coral-isolated *Ostreobium* strains from two distinct *rbc*L lineages, representative of IndoPacific environmental phylotypes, that had been pre-acclimatized (>9months) to three ecologically-relevant reef salinities: 32.9, 35.1 and 40.2 psu. Bacterial phylotypes were visualized at filament scale by CARD-FISH in algal tissue sections, localized to the surface, within filaments or in the algal mucilage. *Ostreobium*-associated communities, characterized by bacterial 16S rRNA metabarcoding of cultured thalli and corresponding supernatants, were structured by host genotype more than salinity and partly overlapped with those of environmental (*Ostreobium*-colonized) coral skeletons. Alphaproteobacteria dominated the thalli communities, enriched in Kiloniellaceae or Rhodospirillaceae depending on algal genotype. A small core microbiota composed of 7 ASVs (∼1.5% of thalli ASVs, 19%-36% cumulated proportions), shared by multiple cultures of both *Ostreobium* genotypes and persistent across 3 salinities, included putative intracellular Amoebophilus and Rickettsiales bacteria. This novel knowledge on the taxonomic diversity of *Ostreobium* bacterial associates paves the way to functional interaction studies within the coral holobiont.

## Introduction

Among reef benthic algae, the siphonous green algae in the genus *Ostreobium* (Chlorophytes) are major carbonate bioerosion agents, dissolving up to a kilogram of reef carbonate per m^2^ per year (Tribollet, 2008). They are ubiquitous colonizers of intertropical, subtropical and temperate reef carbonates, from photic shallow-waters to mesophotic depths (Littler *et al*., 1985; Gonzalez-Zapata *et al*., 2018; Rouzé *et al*., 2021).

Forming an entire Ostreobineae suborder within the Bryopsidales order in the Ulvophyceae class (Sauvage *et al*., 2016; Verbruggen *et al*., 2017), these genetically highly diverse algae are cryptic. Indeed, they are the only Bryopsidales to grow as exclusively microscopic filaments. Their thallus is composed of a network of interwoven, branching, giant coenocytic filaments, 10 to 20 µm in diameter and up to several hundred µm in length, with multiple nuclei and chloroplasts, and cytoplasmic streaming. These filaments erode galleries within the carbonate (bioeroding form), with epilithic 3-dimensional “tufts of filaments” when they emerge at its surface or when isolated in free-living growth form (Sauvage *et al*., 2016; Massé *et al*., 2020; Pasella *et al*., 2022). In contrast, the coenocytic thalli of other siphonous Bryopsidale green algae such as *Bryopsis* and *Codium* (Bryopsidineae) or *Caulerpa* and *Halimeda* (Halimedineae) form macroscopic three-dimensional algal bodies, with morphologically differentiated frond-like axes (sometimes bearing lateral branches and cortical utricles) and basal anchoring rhizoids (Cocquyt *et al*., 2010).

Prevalent in the skeleton of reef-building corals, the Ostreobineae algal filaments are true endoliths, actively penetrating the hard carbonate via chemical means (Tribollet, 2008), and dominating the endolithic microbial communities hosted by living colonies (Ricci *et al*., 2021). Coral species, which differ in their skeletal morphology and growth, can harbor different *Ostreobium* biomass. Fordyce *et al*. (2020) showed a correlation between skeletal density (which impacts light trapping) with biomass and chlorophyll concentration of phototroph endolithic communities dominated by *Ostreobium*. In slow-growing massive corals, their biomass can form dense, visible green bands beneath the host tissue (Lukas, 1974; Le Campion-Alsumard *et al*., 1995). In fast growing branched corals, their networks of filaments are less visible, being “diluted” in the white skeleton mass just below the living coral tissues, and increasingly abundant towards branch bases (Godinot *et al*., 2012; Massé *et al*., 2018). These eroding algae emerge in tissue-free areas for example at the surface of fish bite scars, and after bleaching-induced coral decay and death (Le Campion-Alsumard *et al*., 1995; Tribollet, 2008; Leggat *et al*., 2019). Transmitted horizontally from free-living environmental propagules to the skeleton of juvenile coral recruits, these ectosymbionts were detected as early as 7 days after coral larval metamorphosis and onset of calcification (Massé *et al*., 2018). There is growing consensus that these photoautotroph algal endoliths and their associated prokaryotes have an overlooked functional role in the health of their coral host (reviewed by Pernice *et al*., 2020; Ricci *et al*., 2019; van Oppen and Blackall, 2019): they bloom during coral bleaching episodes, and support bleached coral recovery via suggested provision of alternative photosynthetic carbon nutrients (Schlichter *et al*., 1995, Fine and Loya 2002, Sangsawang *et al*., 2017). Other suggested roles involve contribution to nitrogen cycling (Massé *et al*., 2020; Cárdenas *et al*., 2022) and increased host photoprotection based on recorded modifications of skeletal optical properties (Galindo-Martínez *et al*., 2022).

Investigating the spatially-resolved taxonomic diversity of endoliths within a single coral host colony is an emerging field. Indeed, multiple Ostreobineae genotypes co-exist with diverse prokaryote communities, and the spatial structure of endolithic genetic diversity is patchy, at centimeter scale (Marcelino *et al*., 2017a). *Ostreobium* populations are typically more well-characterized and more genetically diversified in massive slow-growing than branched fast-growing coral host colonies, both in reef (Marcelino *et al*., 2017b) and aquarium settings (Massé *et al*., 2018).

However, the composition of the bacterial communities specifically associated with *Ostreobium* remains a black box, both for isolated strains and within natural coral host colonies. It remains to be determined whether the different *Ostreobium* genotypes are associated with distinct bacterial communities. In a previous study (Massé *et al*., 2020), fatty acid bacterial markers were detected in *Ostreobium* strains isolated from aquarium-propagated *Pocillopora* coral colonies. The diversity of coral-associated endolithic bacterial communities is being captured mostly via amplicon sequencing of variable regions of the bacterial 16S rRNA gene marker from skeletal DNA extracts (Sweet *et al*., 2011; Yang *et al*. 2019, Ricci *et al*., 2022), sometimes in parallel with algal plastid-encoded markers (Marcelino *et al*., 2017a; Marcelino *et al*., 2017b). Whole metagenome-sequencing approaches have seldom been used, restricted to massive slow-growing Hawaiian or Red Sea corals, to quantify the number of genes attributed to bacteria, archaea, viruses, and microeukaryotes from extracted DNA (Wegley *et al*., 2007), recently including *Ostreobium* and fungi (Cárdenas *et al*., 2022). One limitation of 16S rRNA gene based studies is the underestimated retrieved diversity of Ostreobineae, which is better accessed using chloroplast-encoded *rbc*L and *tuf*A gene markers more specific to Bryopsidales algae (Sauvage *et al*., 2016; Marcelino and Verbruggen, 2016; Massé *et al*., 2018). Without documenting thoroughly the genetic diversity of *Ostreobium* within the endolithic assemblages, it is impossible to disentangle the skeletal bacterial fraction specific to *Ostreobium* genotypes from the other bacterial residents selected directly by the host coral genotype or by the environment.

To this day, there is no published survey of the taxonomical composition of the bacterial communities associated with *Ostreobium* and its specific bacterial phycosphere, and no information regarding the influence of environmental factors such as salinity on its composition. Temperature is the most studied abiotic factor that impact the microbiome composition of algal members of the coral holobiont: it is known to modify the bacterial communities of cultured Symbiodiniaceae endosymbionts (Camp *et al*., 2020); recent studies also showed shifts in *Ostreobium* abundance and diversity in response to warming temperatures and ocean acidification (reviewed by Maire *et al*., 2022). However, salinity’s impact on the diversity and physiology of Ostreobineae algae remains unexplored, and their salinity tolerance range is unknown.

Climate change is affecting salinity across oceans, with centennial records showing seawater becoming saltier, for example in the Adriatic Sea (+0.18 psu in the last century; Lipizer *et al*., 2014), and 1950-2000s records of global water cycle intensification, with wet areas getting wetter and less saline, and dry regions saltier (Durack *et al*., 2012). Shallow-water corals thrive in a broad range of salinities in tropical and subtropical areas, from 20 psu in brackish estuaries to 40-42 psu in the Red Sea (Kleypas *et al*., 1999). Locally, and especially in shallow water reef flats, seawater salinity is affected by heavy rainfall and freshwater run-off during episodes of tropical storms, and by evaporation during heat waves of increased frequency and severity. High salinity stress on coral reefs is also caused by brine discharge from desalination plants (Petersen *et al*., 2018), a local pollution by-product from geo-engineering operations to supply increasingly scarce freshwater resources.

High salinity is known to impact the physiology of tropical corals (aquarium-acclimatized to 34-40 psu salinities in Ferrier-Pages *et al*., 1999, or reef-exposed to brine discharge from desalination plant in Petersen *et al*., 2018). Bacterial communities of *Fungia granulosa* corals shifted in response to a 29 days exposure to hypersalinity treatment, with selection of bacterial taxa conferring benefits to the coral host (Röthig *et al*., 2016), but this study did not focus on the skeletal endolithic compartment. Salinity changes impact the community composition of the Symbiodiniaceae dinoflagellates endosymbiotic of coral tissue, which may in turn adjust their bacterial community composition. Indeed, core bacteria specific to Symbiodiniaceae were recently revealed via combined bacterial 16S rDNA metabarcoding of cultured strains and FISH visualization *in hospite* (Lawson *et al*., 2018; Maire *et al*., 2021a). Identifying and visualizing bacterial associates of *Ostreobium* filaments, the endolithic coral algal ectosymbiont, remains an open challenge, which can be addressed by using coral-isolated *Ostreobium* strains. Indeed, such strains represent valuable biological model tools to experimentally investigate this algal biology in simplified *in vitro* systems (Massé *et al*., 2020; Pasella *et al*., 2022).

Here, the diversity of *Ostreobium*-associated bacteria was investigated in two distinct lineages, each exposed to three distinct salinity levels. Experiments were conducted on two genotyped strain representatives of *rbc*L clade P1 (010) and *rbc*L clade P14 (06) lineages *sensu* Massé *et al*., 2020, which both belong to *Ostreobium* lineage 3 *sensu* Tandon *et al*., 2022, within the Ostreobineae suborder. The culture medium salinity was manipulated in order to investigate *Ostreobium* tolerance to salinity increase and related changes in its associated bacteria, in multiple cultures of each strain, long-term acclimatized (9-13 months) to ecologically relevant low (32.9 psu), medium (35.1 psu) and high salinities (40.2 psu).

CARD-FISH experiments on *Ostreobium* thalli acclimatized to low and high salinities (32.9 and 40.2 psu) were performed to visualize at filament scale the bacterial phylotypes in algal tissue sections. Bacterial 16S rRNA gene amplicons were sequenced to characterize the taxonomic diversity of bacteria in cultured *Ostreobium* thalli and corresponding supernatants, according to genotype and salinity, and compared to environmental *Ostreobium*-colonized endolithic communities within coral skeletons.

## Materials and Methods

### Ostreobium *strains and acclimatization to 3 salinities*

The bacterial communities associated to two *Ostreobium* strains, 010 (MNHN-ALCP-2019-873.3) and 06 (MNHN-ALCP-2019-873.2) (Massé *et al*., 2020) from the RBCell ALCP collection of microalgae, Paris, France (Hamlaoui *et al*., 2022) was studied after long-term pre-acclimatization to three salinities: 32.9 psu (13 months), 35.1 psu (12 months) and 40.2 psu (9 months), representative of the natural salinity range in tropical coral reef ecosystems. Both strains were initially co-isolated in 2016 from a healthy branch tip of the same coral host colony *Pocillopora acuta* (Lamarck 1816) propagated in long term, closed-circuit cultures at the Aquarium Tropical, Palais de la Porte Dorée, Paris, France (initially imported from Indonesia before 2010). These *Ostreobium* strains have previously been genotyped to two distinct genetic lineages, i.e. clades P1 for 010 and P14 for 06, based on the *rbc*L taxonomic marker gene (>8% sequence divergence; Massé *et al*., 2020). Free-living growth forms of each *Ostreobium* strain were cultured as suspended tufts of filaments in ventilated T-25 plastic flasks (D. Dutscher, France) containing 25 mL of Provasoli Enriched Seawater medium (PES; Provasoli, 1968; Massé *et al*., 2020). Originally isolated from their coral host in the presence of low doses of penicillin (100 U/ml) and streptomycin (100 µg/mL) antibiotics, they were propagated for 4.5 years, with penicillin and streptomycin (Massé *et al*., 2020) to prevent overgrowth by opportunistic bacteria. Although routinely used in animal cell culture, these generic antibiotics are known to be not efficient against most native marine bacteria (Domart-Coulon and Blanchoud, 2022). And, indeed, bacterial fatty acid markers were consistently detected within chloroform extracts of these strains (Massé *et al*., 2020).

However, to remove antibiotics selection pressure on the algal microbiota, penicillin was omitted for the last 9 months and streptomycin for the last month before sampling for DNA extraction. Long-term *ex-situ* culture introduces microbiota selection pressure but also reduces the variability of abiotic parameters, stabilizing algal phenotypes, for reproducible experimental testing of salinity response under *in vitro* controlled physico-chemical conditions.

For each *Ostreobium* strain and salinity levels, multiple cultures (n=4) were used. For both strains, a parental culture stock (initial algal thallus) was split in two subcultures, each after 6 to 8 months of growth were resplit in 2, resulting in four distinct subcultures per strain (Supplementary Figure 1). Initially grown at 29 psu, all subcultures were gradually acclimatized before the experiment to salinities 32.9 (low salinity, temperate NEAtlantic), then 35.1 (average salinity of Indo-Pacific seawater) and finally to 40.2 psu (Red Sea salinity) by gradually increasing the salinity by about 2-3 psu/month at each monthly medium renewal (Supplementary Figure 1). Artificial seawater of each salinity was prepared by dissolving commercial Reef Crystals sea salt mix (Aquarium Systems, Sarrebourg, France) in H2O (38.6, 40 and 42 g.L-1 for 33.5, 35.5, 40.5 psu, respectively), adjusting pH to ∼8.15 (average seawater pH) then checking final salinity with a CDC401 (HACH) conductivity probe.

Salinities of fresh and spent PES culture medium were controlled at monthly medium renewal, and the measured salinity drift was less than 5% (data not shown). Cultures were incubated at 25°C with 40 rpm orbital shaking (INFORS horizontally rotating platform incubator, Switzerland), under air, and a 12h light / 12h dark cycle of illumination with white fluorescent light intensity of 31 ± 5.5 μmol.m^-2^.s^-1^ (measured with a spherical quantum sensor Li-Cor, USA). Culture medium and T-25 flask were renewed every 4 weeks in all subcultures. Given their long-term (14 – 20 months) physically separate maintenance since sampling from initial stock cultures and before sampling for microbiota characterization, these multiple cultures are assumed to be independent biological replicates. During the year of acclimatization to the three studied salinities, all algal cultures were healthy, with growth and chlorophyll content conserved across the 3 salinity treatments, as measured with visible monthly biomass increase and stable green coloration of the algal tufts of filaments.

### *Catalyzed Reporter Deposition Fluorescence* In Situ *Hybridization (CARD-FISH)*

*Ostreobium* thalli of two cultures of each strain (010, 06), acclimatized either to low or high salinities (32.9 and 40.2 psu), were scalpel-microdissected into fragments (3 mm x 3 mm), allowed to recover for 1 week *in vitro*, then fixed in 4% paraformaldehyde in Sorensen buffer (0.1M) supplemented with sucrose (0.6M) for 2h at room temperature (RT) and then stored at 4°C. Fixed algal thalli (n=4) were rinsed in Sorensen buffer (0.1 M) and embedded in sterile 1.5 % agarose for protection of the fragile network of filaments, before dehydration in an increasing series of ethanol (70% to 100%). Ethanol was substituted overnight by butanol-1 and tissue were embedded in Paraplast Plus paraffin (Leica, France). Histological sections (10 µm) were cut with disposable blades on Leica RM2265 microtome and collected on Superfrost Plus^™^ slides. Sections were deparaffinized with 100% toluene, substituted in ethanol (100% then 96%) and air-dried. They were treated with 3% hydrogen peroxide solution (Sigma) for 30 minutes at RT to inactivate endogenous peroxidases then rinsed three times in Tris-HCl 20 mM pH 8.0. To allow oligonucleotide probe’s access to bacterial 16S rRNA, sections were treated with HCl (10 mM, 10 min at RT), lysozyme (10 mg/mL, 15 min at 37°C) and proteinase K (1 µg/mL, 15 min at 37°C) dissolved in Tris 20 mM, with three rinses in Tris-HCl 20 mM pH 8.0 after each step. *Ostreobium* sections were hybridized using a EUB338mix of three probes: EUB338 (5’ GCTGCCTCCCGTAGGAGT-3) (Amann *et al*., 1990), EUB338 II (5’GCAGCCACCCGTAGGTGT 3′) and EUB338 III (5′ GCTGCCACCCGTAGGTGT 3′) (Daims *et al*., 1999) (v/v/v 2:1:1) a combination which detects most bacteria. Negative controls were performed using non-EUB probe (5’ CTCCTACGGGAGGCAGC 3′) (Amann *et al*., 1990) on serial sections of the same paraffin block, mounted on the same slide. We checked that chloroplast encoded 16S rRNA sequences (from published *Ostreobium* genomes) did not align (3 mismatches) with reverse sequences of EUB probes, excluding artefactual probe hybridization on chloroplasts. To amplify the hybridization signal, we used horseradish peroxidase - conjugated oligonucleotidic probes (HRP-probes, purchased from http://www.biomers.net/, Germany) diluted at 5 ng/µl in hybridization buffer (HB), composed of 0.9 M NaCl, 20 mM Tris-HCl pH 8.0, 0.01% SDS, 10% dextran sulfate, 1% blocking reagent (Roche Diagnostics, Mannheim, Germany), and 35% (v/v) formamide (for both EUB338 mix and nonEUB). Each section was covered with ∼10µL of HRP-probe in hybridization buffer (exact volume adjusted to cover the surface of each section). Hybridization was performed in wet slide chamber at 42°C for 2h30, then stabilized at 43°C for 15 min in washing buffer (5 mM EDTA, 20 mM Tris-HCl pH 8.0, 0.05% SDS, 42mM NaCl and 35% (v/v) formamide), followed by two steps of rinsing in PBS 10 mM pH 7.4 at RT. The HRP-probe signal was then amplified for 15 min at 37°C in amplification buffer (PBS 0.8X pH 7.4, 1.6M NaCl, 8% dextran sulfate, 0.08% blocking reagent) containing 10 µL H2O2 0.3% and 2.5 µL of Alexa488 fluorochrome-labeled tyramides (Invitrogen). Sections were washed two times in PBS for 10 min at RT, rinsed for 1 min with milliQ water, air-dried and mounted in Fluoroshield containing DAPI (Sigma).

Thalli sections (10µm) were observed with a Zeiss AxioZoom V16 macroscope equipped with a X2.3 Plan NeoFluar objective (NA 0.57, FWD 10.6mm), HXP200C metal halide lamp and Zeiss AxioCam 506 camera. Fluorescence images were acquired in single plane or z-stack series (8-12µm depth, step 1µm) and same exposure settings for treatment (EUBmix) and control (nonEUB) with ZenBlue 3.2 software at the MNHN photonic microscopy platform (CeMIM). Alexa488 positive bacterial morphotypes were visualized in Extended Depth of Focus projection (Wavelets method) of deconvoluted (nearest neighbor) z-stack series of images acquired at Ex 450-490 nm (Em 500-550 nm). Dapi-stained DNA was visualized at Ex 340-390 nm (Em 420-470 nm). Multichannel images were assembled in ImageJ (Fiji), with similar visualization settings (intensity and contrast) for treatment (EUBmix) and controls (non EUB). Multiple algal tissue areas in 1 to 2 serial sections per treatment or controls were imaged per slide, with 1-2 slides imaged per culture.

### Sampling and DNA extraction

Biomass (thalli) of *Ostreobium* (n=24, i.e. 4 biological replicates x 3 salinities x 2 strains, each with dry weight 12.55 ± 4.54 mg) were sampled 9 days after transferring the cultures to fresh PES medium. Tufts of filaments were rinsed three times in ∼20 mL of 0.2µm filter-sterilized artificial seawater (Reef Crystal) at corresponding salinity levels and then lyophilized. The corresponding supernatants (n=24) had been sampled previously at the last transfer (∼25 mL each, after 1 month of *Ostreobium* culture) and filtered through a 0.22 µm GTTP Isopore^™^ membrane filter, stored at -80°C to access the bacterial community composition of strain supernatants. Total DNA extraction of algal biomass and supernatants (i.e. filters) was performed in sterile conditions (under laminar flow hood) using DNeasy PowerSoil™ Kit (Qiagen Laboratories, CA). For *Ostreobium* biomass, 2% polyvinylpyrrolidone (PVP) was added in the lysis buffer to remov m e phenolic compounds. For all DNA extracts, glycogen was added as a nucleic acid carrier (30 µg/ml final concentration) before DNA precipitation. As internal controls, DNA extractions were also performed on fresh PES medium for each salinity (in duplicate, n=6, i.e. 3 salinities x2), 2% PVP (n=1) and a membrane filter (n=1).

DNA extractions from skeleton fragments of environmental *Pocillopora* sp. corals collected at a medium-low salinity reef site, 34.4 psu (Guam, Pacific, n=4), and a high salinity reef site, ∼40.6 psu (Eilat, Red Sea, n=4), were also used as controls for wild populations of *Ostreobium* and associated bacteria (complex coral endolithic assemblages, with microboring Ostreobineae presence confirmed by *tuf*A and *rbc*L amplicon sequencing, see below). Skeletal DNA was extracted after removal of coral tissues, using Quiagen DNeasy PowerSoil™ Kit after protocol from Massé *et al*., 2018. For each reef and salinity level, multiple colonies (n=4) were sampled from the same site. Skeletons of *Pocillopora sp. cf meandrina* corals from Guam island site 2 (13.42 N 144.64 E) are part of the Tara Pacific 2016-2018 collection (M152C01/CO-1008013, M152C02/CO-1008014, M152C05/CO-1008017, M152C06/CO-1008018) with metadata-provided salinity. Skeletons of *Pocillopora* sp. cf *verrucosa* colonies from Eilat (P1, P2, P3, P4) were collected at depth 6-10m on July 4 2019 on the InterUniversity Institute (IUI) reef nursery table (34.92 N 29.50 E) during the European Union’s Horizon2020 ASSEMBLE Plus (grant N°730984) 2018 Transnational Access project CORALBOUCLE-CLIM (collection permit 209/42300 Israel Nature and Parks Authority to M. Fine), with salinity retrieved from the Israel National Monitoring Program at the Gulf of Eilat datasets.

### PCR amplification and Illumina sequencing

The V5-V7 region of the bacterial 16S rDNA (∼400 bp) was amplified using specific primer pair 799F (5’ AACMGGATTAGATACCCKG 3’) and 1193R (5’ ACGTCATCCCCACCTTCC 3’) to limit co-amplification of chloroplastic 16S rDNA of algal host (Vieira *et al*., 2016; Tourneroche *et al*., 2020). Illumina adapter sequence were added to the 5’ – end of the forward and reverse primers. To increase the recovered diversity, PCR amplifications were performed in triplicate for each sample, then pooled during the purification step (see below). Amplification reactions were performed in 25 µL volume containing 2 µL DNA extract template, 0.5 µM of each primer, 2 µL MgCl2 (25mM), 0.5 µL DNTPmix (10 mM), 5 µL Go Taq Flexi Buffer, 0.125 µL of 5X GoTaq^®^ G2 Flexi DNA Polymerase (Promega) in sterile water. The PCR cycling conditions were: 3 min at 95°C, 35 cycles of [20 s at 95°C, 30 s at 54°C, 30 s at 72°C], and 10 min terminal extension at 72°C. Amplified fragments were visualized in 1% agarose gels with SYBRSafe and amplicons from 3 independent positive PCR reactions were pooled, then column purified (Nucleospin^®^ gel and PCR clean-up kit, Macherey-Nagel, France) in EB elution buffer (Qiagen). Amplicons from negative PCR controls were also pooled and purified as internal control (n=1). All purified amplicons (25 µL) were submitted for MiSeq sequencing (Illumina, paired-end, 2×300 bp) to Eurofins Genomics (Germany).

*Ostreobium* was detected in coral skeletons of *Pocillopora* sp. from Guam (Pacific) and Eilat (Red Sea) as previously published for DNA extraction (Massé *et al*., 2018), and amplicon-sequencing of the *rbc*L gene fragment with the primers *rbc*LF250 ([5’ GATATTGARCCTGTTGTTG GTGAAGA 3’] ; modified from Gutner-Hoch and Fine, 2011) and *rbc*L1391R (Verbruggen *et al*., 2009), or the *tuf*A elongation factor gene fragment with primer pair: Oq-tuf (Marcelino and Verbruggen, 2016) / *tuf*AR (Famà *et al*., 2002) (∼490 bp) and *tu*bryoF / *tu*bryoR (∼600 bp, Sauvage *et al*., 2016). Additionally, the *tuf*A marker was amplified from DNA extracts of thalli of *Ostreobium* strain 06 and 010 with both primer pairs. Amplicons were submitted for MiSeq sequencing (Illumina, paired-end) to the MNHN SSM (Service de Systématique Moléculaire) platform (MNHN-CNRS UAR2700 ‘Acquisition et Analyse de Données pour l’Histoire naturelle’).

### Sequence dataset analysis

For bacteria, primer sequences were removed in raw reads using CUTADAPT program (v2.5, Martin, 2011). Paired 16S rDNA amplicon reads were then assembled using FLASH algorithm (2.2.00, Magoc and Salzberg, 2011) with a minimum overlap size of 10 bp. Sequences analysis was performed using QIIME 2 2022.2 (Bolyen *et al*., 2019). Consensus sequences were demultiplexed using DEMUX algorithm. DADA2 plugin was used to remove chimeras, trim sequences to 368 bp, denoise the dataset, and assign reads to Amplicon Sequence Variants (ASVs). ASVs were assigned to taxonomic groups using the feature-classifier plugin and classify-sklearn module in SILVA v138 SSU rRNA database (released December 16, 2019). ASVs affiliated to Eukaryota, Chloroplast, Mitochondria, and Unassigned ASVs (i.e. assigned to none of the three domains of life) were removed from the dataset, as well as all ASVs of the internal controls (ASVs detected either exclusively in fresh PES medium, PVP extraction supplement, membrane filters, and PCR controls, or shared with at least one algal or coral sample, without taking into account relative abundances thresholds). This stringent approach may underestimate natural bacterial diversity, but makes sure that the obtained diversity was not an artifact of contamination. In *Ostreobium* biomass samples, bacterial sequences unassigned below the class level in SILVA v138 were then aligned by BLASTn with 16S RNA-encoding sequences available in GenBank database (consulted on July 21-22 2022): the order could be identified when the unassigned sequence had a similarity of more than 90% with the Genbank reference sequence. Rarefaction curves were performed on rarefied ASV table using QIIME 2 2022.2 (1,080 sequences (minimum library size) sub-sampled for each sample).

Alpha diversity (ASV richness, Shannon and Pielou’s eveness indices) was analyzed on the unrarefied dataset in R software v4.0.5 (package vegan). Differences between groups (salinity treatment and *Ostreobium* genotypes) were tested using the non-parametric Kruskal Wallis test, followed by a pairwise post-hoc analysis of Mann and Whitney with a threshold value of 5% (α=0.05). For beta-diversity, a matrix of Bray-Curtis distances was calculated from the arcsine square root transformed unrarefied bacterial ASV proportions. Variance levels were first compared between environmental coral skeletons *versus in vitro* algal cultures (*Ostreobium* thalli and supernatants) as there was a difference of dataset size (2 salinities studied for environmental corals *versus* 3 for *Ostreobium* cultures). Then, the bacterial communities were compared between algal categories (*Ostreobium* thalli *versus* supernatant), genetic lineages (010 clade P1 *versus* 06 clade P14), salinities (32.9 *versus* 35.1 *versus* 40.2 psu) and their interactions, using permutational multivariate analysis of variance implemented in the ‘adonis2’ function of the vegan package in R (n=999 permutations, α = 0.05) followed by pairwise comparisons (‘pairwise.adonis’ function) with Bonferroni correction.

To plot the bacterial variation among sample categories, Principal Component Analysis (PCA) was performed in R software (v4.0.5) on arcsine square root transformed bacterial ASV proportions (unrarefied data) and compared to Principal Coordinates Analysis (PCoA) of the Bray-Curtis distances. Next, to focus on the microbiota of *Ostreobium* thalli, another PCA was performed to plot the variance structure among algal genetic lineages and salinities, and highlight bacterial ASVs that explain (correlation circle) most of the variance in our datasets (additional PCA at bacterial order level was also performed to visualize thalli community structure at lower taxonomic resolution). Sparse Partial Least Square Discriminant Analysis (sPLS-DA) was used to classify by host lineage or salinity the bacterial community profiles assessed at ASV level (R v3.6.2, package mixOmics, arcsine square root transformed unrarefied proportions, with analysis limited to the 1000 most discriminant ASVs of the high-dimension dataset). To select informative ASV variables, that predict community projection on the Principal Component dimension that best discriminates between factors, we analyzed the correlation of sample projection coordinates with ASV composition. We ranked bacterial ASVs with correlation coefficient >0.7 towards the sample projection on Component 1 of the PLS models. This allowed to identify bacterial ASVs that contribute the most to the classification of groups, and highlighted the most discriminating bacterial ASVs within each model. Subsequent heatmaps illustrated fold changes of these differentially expressed variables among replicates and treatments (R v4.0.5, package “pheatmap”). Venn diagrams were built to identify and visualize the ASVs that are part of the *Ostreobium* core microbiota, the criterion being that these ASVs should be shared by both genetic lineages and occur across all 3 salinities. Data were visualized with R v4.0.5 packages “ggplot2” and “tidyverse”.

For *Ostreobium*, paired-end reads of *tuf*A were assembled and analyzed using an in-house bioinformatics pipeline, which consist of several steps. Primers were removed from the raw reads using CUTADAPT and DADA2 was used for quality filtering and identification of ASVs (Callahan et *al*., 2016). A table of ASVs across different samples was generated. Plastid sequences were then aligned by local BLASTn with *tuf*A -encoding sequences available in GenBank database (consulted 2 May 2022) for classification to *Ostreobium* families *sensu* Sauvage *et al*., 2016.

### Data availability

Sequences generated via Illumina Miseq sequencing during this study were deposited as NCBI project archive “BioProject: PRJNA896951”. Raw bacterial ASV tables (read counts of unfiltered and filtered data), fasta sequences and corresponding taxonomy, as well as fasta sequences of Ostreobineae *tuf*A ASVs detected in coral skeletons has been deposited at figshare with doi: 10.6084/m9.figshare.21953075 (Supplementary Table 1).

## Results

### In situ *localization of bacteria associated to cultured* Ostreobium *filaments*

Bacterial phylotypes revealed by CARD-FISH experiments using universal bacterial probe EUBmix (EUB338 + EUBII + EUB III) hybridized to 16S rRNA were localized within or in close proximity to *Ostreobium* filaments, for both genetic lineages, and at low (32.9 psu) and high (40.2 psu) salinities (Figure 1A, 1B, 1C, 1D). Reconstructions in 3D of z-stacked series of observations showed EUB-positive signal at the surface of *Ostreobium* filaments, indicative of epiphytic bacteria, and EUB-positive signal was also recorded within the filaments, attributed to endophytic bacteria (with uncertainty due to epifluorescence imaging low resolution). Indeed, controls using non-EUB probe were negative (Figure 1A*, 1B*, 1C*, 1D*). Positive CARD-FISH signals also revealed bacteria inside the mucilage between *Ostreobium* filaments (Figure 1). Based on hybridization signal intensity, which relates to ribosomal RNA content, the detected bacteria were metabolically active. Algal nuclei counterstained with DAPI were heterogeneously distributed along the *Ostreobium* filaments (Figure 1), which siphoneous cytological organization revealed: empty portions alternating with areas of several elongated nuclei, concentrated near branching nodes or spread along the filament, together with discoidal chloroplasts. Chlorophyll autofluorescence was not detected here due to ethanol depigmentation before paraffin-embedding, but observed in native filaments. Bacterial abundances were variable depending on algal structures, locally concentrated at the surface of filament envelopes and in the mucilage (especially abundant at high salinity); however, the presence/absence results are qualitative and did not allow testing of a clear differential distribution pattern between structures and cultures.

**Figure 1:**
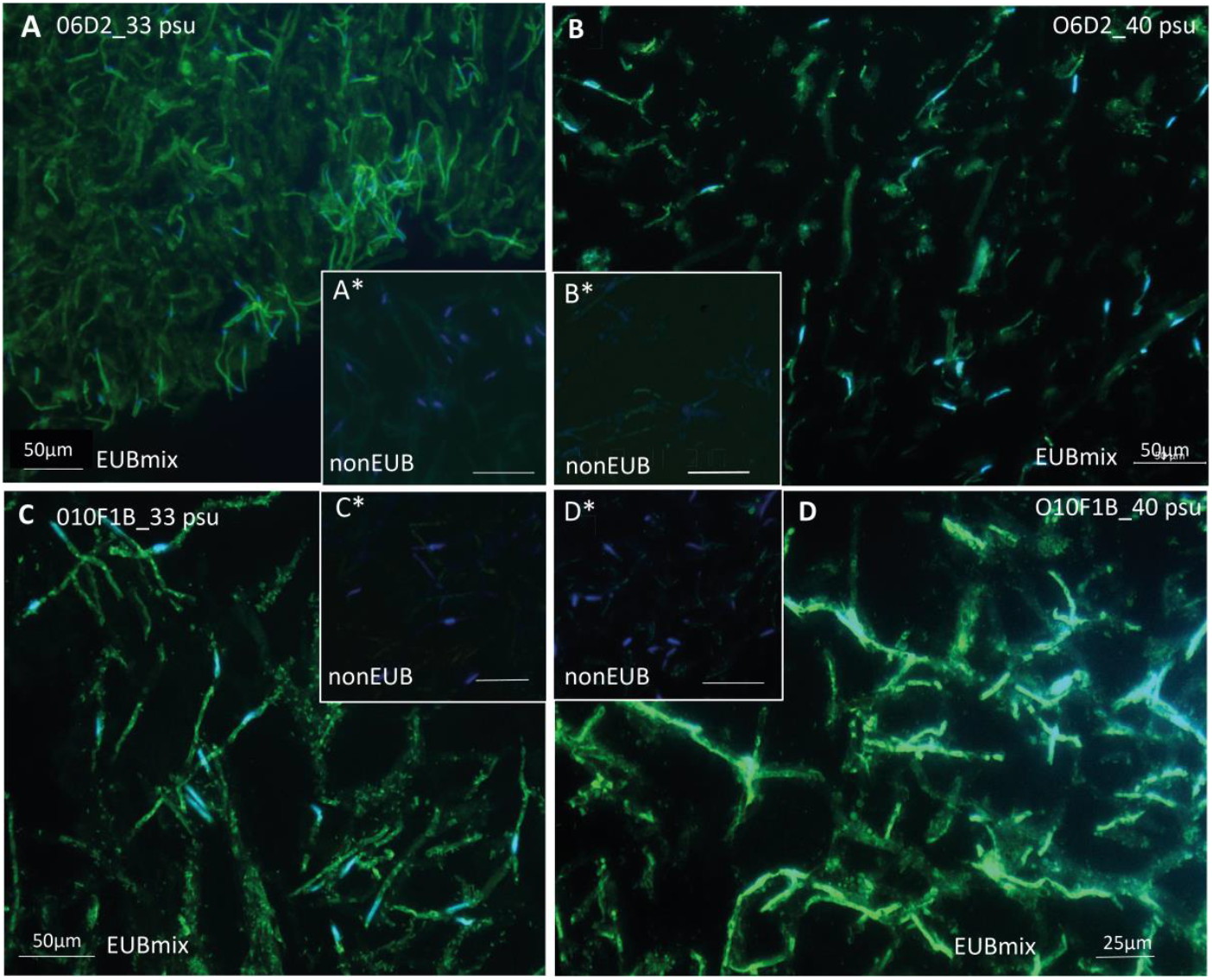
Visualization of bacteria in *Ostreobium* algal tissue sections via fluorescence *in situ* hybridization. Positive hybridization signal of universal bacterial probe mix (EUBmix) detected with Alexa488 (in green, cocci and rod-like morphotypes) on the surface and within *Ostreobium* filaments, or in the mucilage between filaments in deparaffinized sections (10µm) of strains 06 (A, B) and 010 (C, D) cultured at 32.9 psu (A, C) and 40.2 psu (B, D) salinities. Negative controls are hybridized with nonEUB probe (A*, B*, C*, D*). Algal nuclei counterstained with DAPI (in blue) are heterogeneously distributed within the coenocytic thalli, with multiple nuclei containing filaments alongside “empty” filaments (uneven Dapi staining in A and D). Alexa488 positive bacterial phylotypes are tightly associated to filament envelopes (sheaths), with a few intra-filament spots distinct from nuclei and chloroplasts. Positive Alexa488 signals also highlight cluster-aggregated bacterial phylotypes between algal filaments.

### Ostreobium *diversity in environmental* Pocillopora *sp. coral skeletons*

*Ostreobium* detection was confirmed in *Pocillopora* coral skeletons from Guam (Pacific; 34.4 psu) and Eilat (Red Sea; ∼40.6 psu) via *rbc*L and *tuf*A sequence analysis, summarized in Supplementary Table 2. In *Pocillopora* from Guam, *Ostreobium* was detected in all four colony replicates and all ASVs were affiliated to the Odoaceae family based on *tuf*A marker sequences (*sensu* Sauvage *et al*., 2016; >85% sequence similarity to the designated “type” strain). In *Pocillopora* from Eilat, *Ostreobium* ASVs were detected using the *tuf*A marker in 3 out of 4 colonies (no data with *rbc*L marker). In one coral colony, two *Ostreobium* families were detected, Maedaceae and Odoaceae (*sensu* Sauvage *et al*., 2016), as well as ASVs unclassified at family level (nucleotid sequence divergence >15%). In two other coral colonies, ASVs detected were affiliated either to the Odoaceae family or unclassified at family level.

In *Ostreobium* strains, *tuf*A sequences were amplified using both molecular markers but only the *tu*bryoF/*tu*bryoR primers allowed to retrieve *tuf*A sequences of Ostreobineae. For strain 06, one ASV was affiliated to the family Hamidaceae; *sensu* Sauvage *et al*., 2016. For strain 010, all retrieved *tuf*A sequences classified to non-targeted bacterial taxa and our protocol failed to amplify Ostreobineae *tuf*A sequences; although *tuf*A family-level classification of this strain was not possible, previous *rbc*L-based sequencing had however affiliated this strain to the Ostreobineae suborder (Massé *et al*., 2020).

### Quality of Illumina sequencing targeting bacterial16S rRNA V5-V7 gene fragment

A total of 8,247,382 sequences of the V5-V7 region of 16S rDNA were obtained from the 65 samples using Illumina MiSeq sequencing. After chimera detection and quality filtering, between 52 and 89% of the sequences were retained from each sample. These represented a total of 3,930 distinct bacterial ASVs (unrarefied dataset) after removal of ASVs affiliated to Eukaryota (2 ASVs), Chloroplast (4 ASVs), Mitochondria (7 ASVs), Unassigned sequences (11 ASVs) and removal of internal controls (all 1,411 ASVs exclusively detected in controls, and the 222 ASVs shared between controls and at least one sample, no matter their relative abundances). Rarefaction curves of *Ostreobium* thalli and their corresponding supernatants (Supplementary Figure 2, rarefied ASV table) reached a plateau indicating that a reasonable sequencing depth had been attained. Rarefaction curves of environmental corals tended to flatten out (Supplementary Figure 2) suggesting that additional rare bacterial taxa are probably present but undetected in these skeletal samples.

### *Diversity and structure of bacterial communities in cultured* Ostreobium versus *coral skeletons*

Alpha-Diversity indices measured at high taxonomic resolution (ASVs) on unrarefied data for *Ostreobium* algal thalli, their corresponding culture supernatants, and environmental *Pocillopora* sp. corals (endolithic assemblages) are presented in Figure 2 as a function of genotype and salinity (individual values of ASV richness and Pielou’s evenness indices for each sample are provided in Supplementary Table 3). Between 24 and 569 bacterial ASVs were detected in individual samples. Cumulated ASVs, specific or shared between categories (thalli / supernatants / coral skeletons) are illustrated in Venn diagrams (Figure 3A).

**Figure 2:**
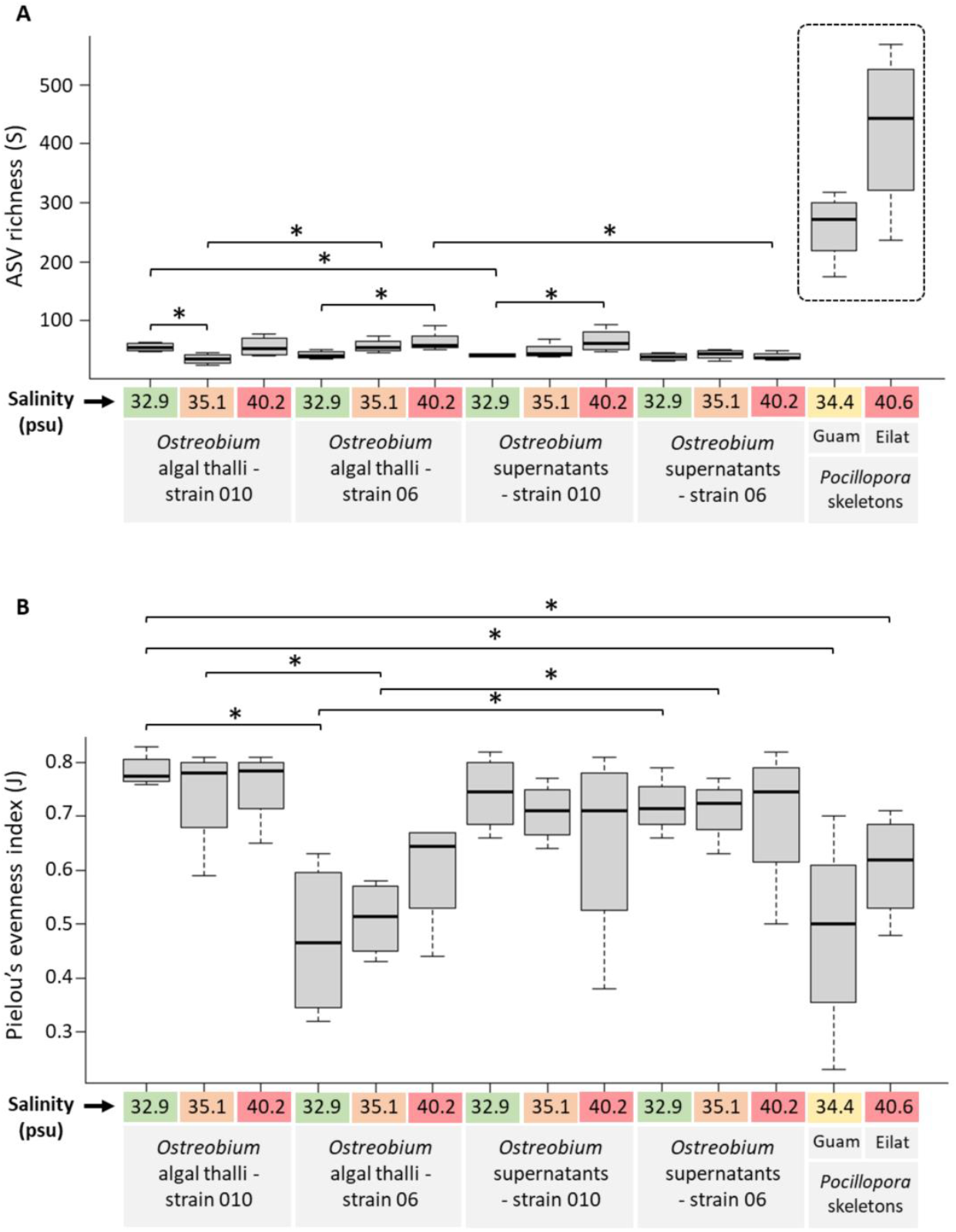
Alpha diversity of the bacterial communities (ASVs) of cultured *Ostreobium* thalli and supernatants (depending on algal genotype and salinity), and of environmental *Pocillopora* coral skeletons. (A) ASV richness and (B) Pielou’s evenness index were calculated on unrarefied ASV data. * significant differences (Kruskal-Wallis, with pairwise post-hoc Mann & Whitney test, p<0.05) between comparisons of interest (e.g. between genotypes for the same salinity, between salinities for the same genotype, between a thallus at a given salinity and its corresponding supernatant, and between cultured *Ostreobium* thalli / supernatants and coral skeletons). Dotted box in (A): Guam and Eilat environmental corals displaying significantly higher alpha-diversity than all cultured *Ostreobium* samples (thalli and supernatants whatever genotype and salinity).

**Figure 3:**
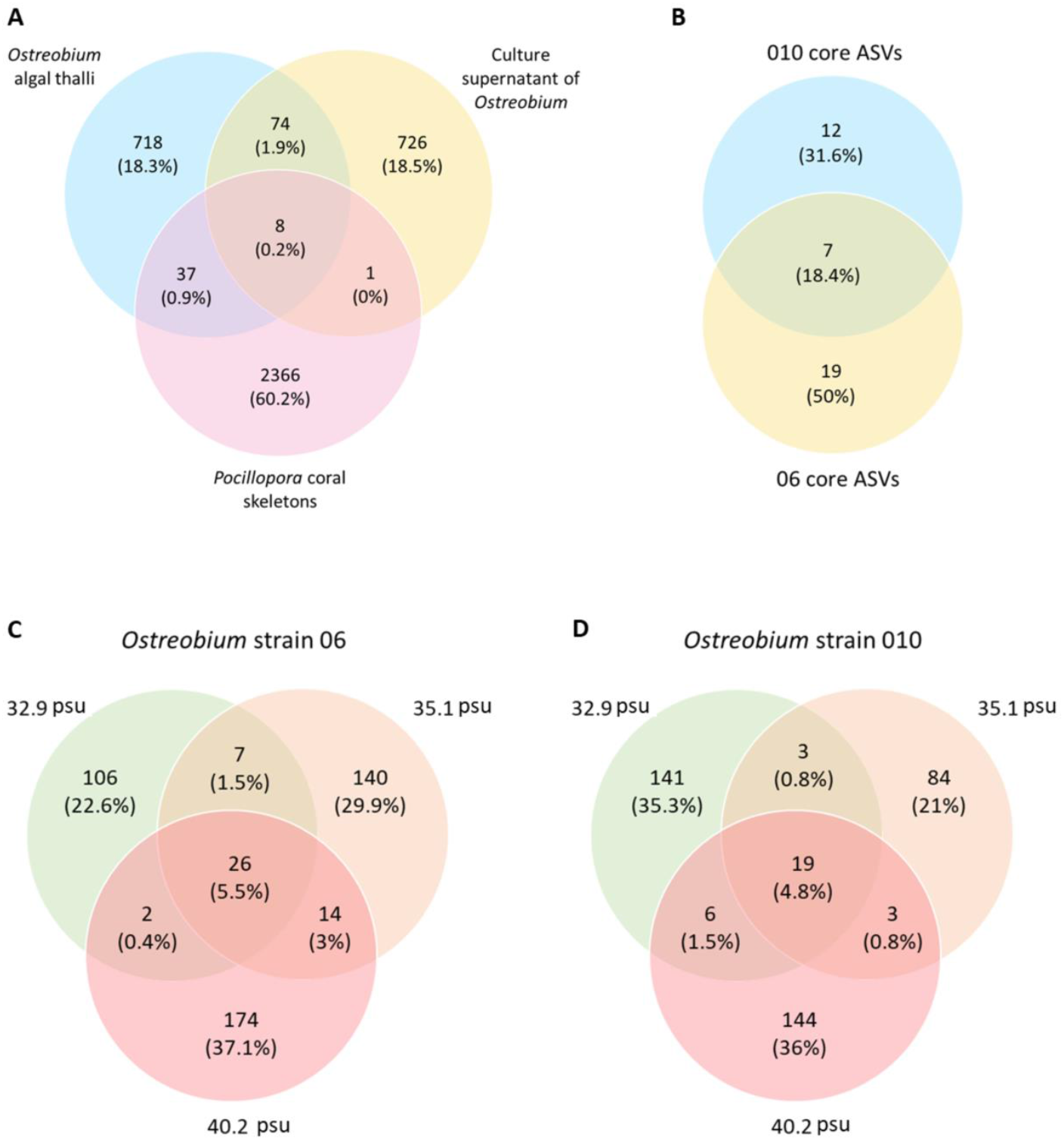
Venn diagrams of bacterial ASVs distribution (A) across cultured *Ostreobium* thalli and supernatants, and environmental *Pocillopora* coral skeletons. (B) Core microbiota of *Ostreobium* shared by both algal genotypes: 010 *rbc*L clade P1, 06 *rbc*L clade P14. Only ASVs conserved across 3 salinities for each genotype are represented (detailed in Supplementary Table 5). (C, D) Bacterial ASVs distribution across salinities (32.9, 35.1 and 40.2 psu) for *Ostreobium* strains (C) 06 and (D) 010.

In cultured algal thalli, a total of 837 cumulated ASVs were detected from both *Ostreobium* genotypes at all 3 salinities (total n=24 cultures; Figure 3A) with totals of 469 and 400 ASVs in the biomass of strains 06 (n=12; Figure 3C) and 010 (n=12; Figure 3D), respectively (similar average 54±16 ASVs per sample for 06, and 49±15 ASVs for 010; Figure 2A and Supplementary Table 3). A total of 809 cumulated ASVs were detected in supernatants from both genotypes at all 3 salinities (n=24, Figure 3A). Between 5.7 and 8.8% of bacterial ASVs were shared between *Ostreobium* thalli of a genotype at a given salinity and its corresponding supernatants.

In environmental corals, i.e. complex endolithic assemblages of *Pocillopora* sp. from Guam (Pacific) and Eilat (Red Sea) including diverse Ostreobineae phylotypes (see above and Supplementary Table 2), a total of 2,412 cumulated bacterial ASVs were detected (total n=8 skeletons, Figure 3A; average 259±61 per sample for Guam and 423±141 for Eilat). Thus, the bacterial ASVs richness detected within “wild” *Ostreobium*-colonized *Pocillopora* skeletons from Guam and Eilat was 5-8 fold higher than within cultured *Ostreobium* thalli (both 010 and 06; Figure 2A). A total of 45 bacterial ASVs were shared between *Ostreobium* thalli and environmental corals (5.4% of thalli’s ASVs, 1.8% of coral ASVs; Figure 3A), indicating partial overlap of bacterial diversity between cultured and “wild” *Ostreobium* (also visualized by PCA analyses of community structures, see section below). In contrast, only 9 ASVs (less than 0.4% of coral ASVs) were shared with *Ostreobium* culture supernatants (Figure 3A).

Alpha-diversity equitability (Pielou’s evenness) in cultured *Ostreobium* algal thalli of strain 010 was stable across salinities and higher than in strain 06, with average values of 0.79±0.03 (n=4), 0.74±0.10 (n=4) and 0.76±0.07 (n=4) for salinities 32.9, 35.1 and 40.2 psu, respectively (Figure 2B). In *Ostreobium* thalli of strain 06, Pielou’s evenness was lower but increased with salinity from 0.47±0.15 (n=4) to 0.51±0.07 (n=4) and then to 0.60±0.11 (n=4) at 32.9, 35.1 and 40.2 psu, respectively. Among thalli, Pielou’s evenness index was significantly different between genetic lineages (strains) at 32.9 and 35.1 psu (Figure 2B; p=0.029), but not at 40.2 psu (p=0.11). Overall, these results indicate high differentiation and stable, even distribution of microbiota across salinities for thalli of strain 010. A pattern towards diversification (increased Shannon index, Supplementary Table 3) and homogeneization (higher Pielou’s evenness index) at highest salinity (40.2 psu) was detected for thalli of strain 06. Among culture supernatants, Pielou’s evenness was stable whatever the salinity, with average values of 0.70±0.12 (n=12) for 010 and 0.71±0.08 (n=12) for 06. In contrast, within coral skeletons’ complex endolithic assemblages, the Pielou’s evenness index was lower and highly variable across biological replicates, with average values of 0.48±0.19 in *Pocillopora* sampled at Guam (Pacific) and 0.61±0.10 in *Pocillopora* from Eilat (Red Sea) (Figure 2B and Supplementary Table 3).

The structure of bacterial communities associated to cultured *Ostreobium* thalli, their supernatants, and *Pocillopora* skeletons was visualized by Principal Component Analysis at ASV (Figure 4A) or order (Supplementary Figure 3A) levels of taxonomic resolution. There was a great dispersion of microbiota profiles of cultured *Ostreobium* thalli, partly overlapping with corresponding supernatants, and also overlapping with profiles of environmental corals when projected on Component 1 (Figure 4A; 11%, PC1). Bacterial assemblages of cultured thalli and environmental corals projected however separately on Component 2 (Figure 4A; 7%, PC2). Similar partial overlap of bacterial profiles between cultured *Ostreobium* (thalli and supernatants) and coral skeletons was also visualized by Principal Coordinate projections of Bray-Curtis distances measured at ASV level (data not shown). Differences between cultured and environmental categories were statistically significant (pairwise “adonis” comparisons after permutational multivariate analysis of variance, F=3.04, df1, p=0.001; Supplementary Table 4A). In *Ostreobium* cultures, there was no significant difference between communities of supernatants and corresponding thalli (F=1.012, df1, p=0.425; Supplementary Table 4B) but differences were significant between thalli genotypes (strain 06 : *rbc*L clade P14 *versus* strain 010: *rbc*L clade P1, F=3.873, df1, p=0.001), between salinities (pairwise comparisons, p=0.001, F values reported in Supplementary Table 4C) and the interaction of algal genotype x salinity (F=2.413, df1, p=0.003): each genotype had a specific bacterial profile which responded differently to salinity increase.

**Figure 4:**
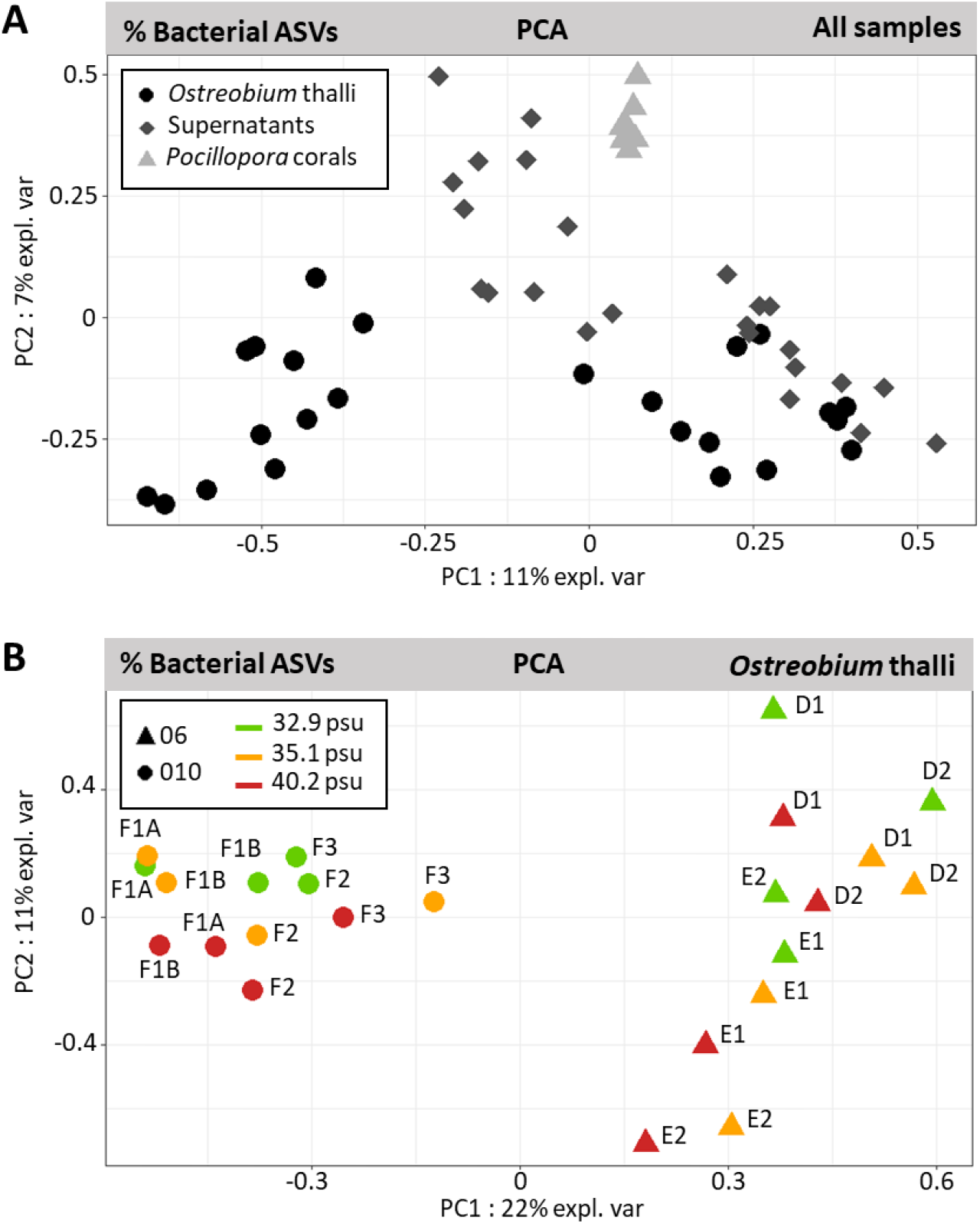
Structure of the bacterial communities visualized by Principal Component Analysis at ASV level (arcsine square root transformed of the unrarefied proportions data). (A) Partial overlap between communities of cultured *Ostreobium* thalli, their supernatants and those of coral skeletons. (B) Clustering of *Ostreobium* thalli bacterial community profiles by algal genetic lineage (010: *rbc*L clade P1, 06: *rbc*L clade P14) more than salinity. Culture code names (FXX, EXX, DXX) are indicated in the score plot.

Within cultured *Ostreobium* thalli, PCA visualization of the variance of bacterial communities among genotypes and salinities confirmed a clear separation according to algal genotype on Principal Component 1 (at ASV level, Figure 4B, 22%, PC1), without clear trend for the salinity factor. Correlation circle analysis of PCA (at order level, Supplementary Figure 3) showed that the clustering of bacterial profiles by algal genotype was explained by 2 orders within the Alphaproteobacteria, namely Kiloniellales in strain 010, *versus* Rhodospirillales in strain 06. At ASV level, the bacterial Rhodospirillaceae ASV16 explained most of the separation between algal genotypes (correlation circle plot of PCA at ASV level, Supplementary Figure 4B). Both Rhodospirillales and Kiloniellales (output from SILVA v138) have since been re-classified as distinct family level lineages (Rhodospirillaceae and Kiloniellaceae) in the class Rhodospirillales (NCBI taxonomy browser, Degli Esposti *et al*., 2019).

### *Taxonomic composition: identification of a core* Ostreobium *bacterial microbiota*

Barplot distribution of bacteria relative abundances in cultured *Ostreobium* thalli, their supernatants, and *Pocillopora* coral skeletons are illustrated at order level in Figure 5 (also at class level, Supplementary Figure 5; taxonomy from SILVA v138 December 16, 2019). A total of 135 bacterial orders (51 classes) were detected, of which 95 (34 classes) had relative abundances below 1%. High inter-individual variability was observed at order level (Figure 5; also at class level: Supplementary Figure 5) among genotypes, salinities and coral skeletons. The Alphaproteobacteria was the most abundant class detected across all samples (15.6-94.2% cumulated abundances), represented by 20 orders (8 < 1%). Within the Alphaproteobacteria, the most abundant order in *Ostreobium* thalli of strain 010 (clade P1) was the Kiloniellales (33.3±16%, n=12) detected in all samples. This order was also detected in all corresponding culture supernatants with similar relative abundances of 31±18.9% (n=12), and in all *Pocillopora* coral skeletons but at 6-fold lower relative abundances of 5.1±6.4% (n=8). In contrast, in *Ostreobium* thalli of strain 06 (clade P14), the most abundant and prevalent alphaproteobacterial order was the Rhodospirillales, with average relative abundances of 36.8±19.9% (n=12, i.e. all thalli). The Rhodospirillales were however much less abundant in corresponding supernatants (2.9±2.3%, detected in 9/12 samples) and in *Pocillopora* coral skeletons (1.0±2.4%, detected in 6/8 samples).

**Figure 5:**
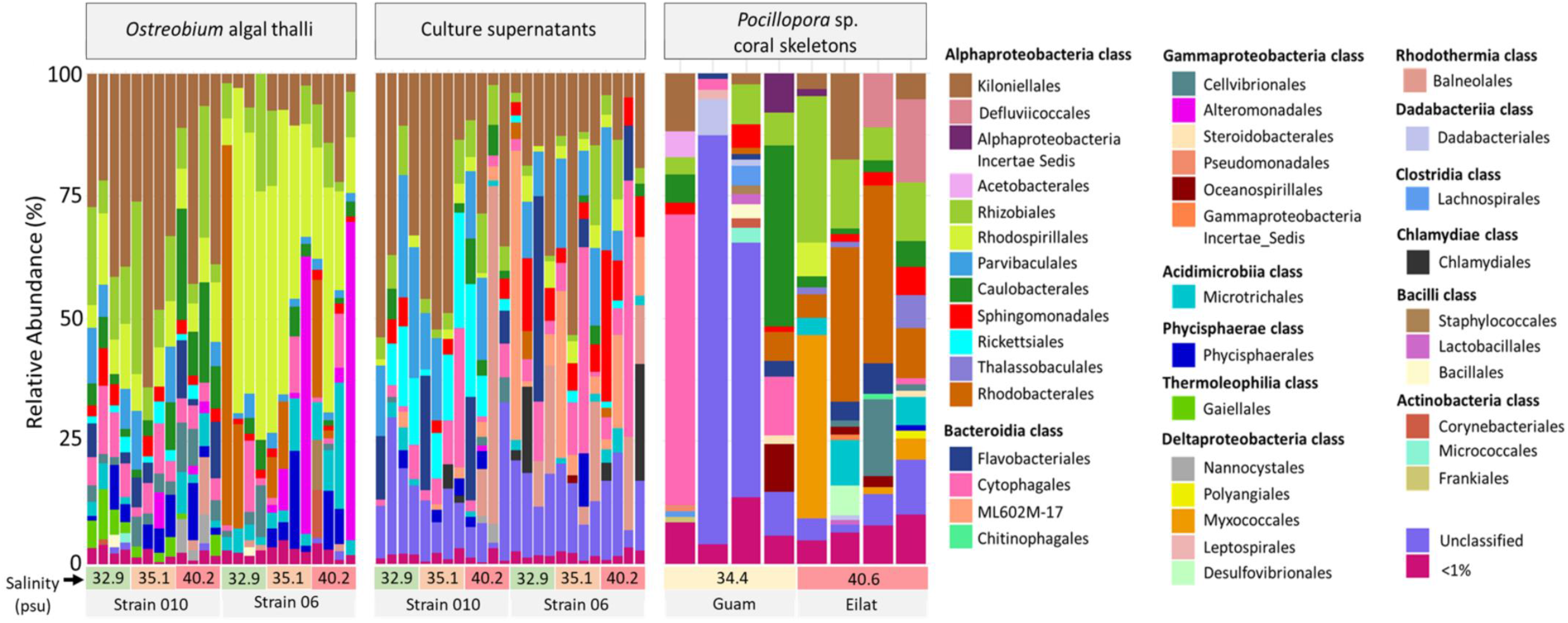
Taxonomic composition of bacterial orders in cultured *Ostreobium* thalli and supernatants, and the environmental *Pocillopora* coral skeletons. Salinity and algal genotype or reef site are indicated below each individual sample column. Orders detected at trace levels are aggregated in the <1% abundance category.

The Bacteroidia class was detected in all samples of *Ostreobium* thalli and culture supernatants but with higher abundance in supernatants (21±19.4%, n=24) than thalli (7.0±5.6%, n=24). This class was also detected in all coral skeletons, at highly variable abundance (12.1±19.8%, n=8).

Gammaproteobacteria, Acidimicrobiia and Phycisphaerae classes were more abundant in *Ostreobium* thalli than corresponding supernatants. Gammaproteobacteria were detected in all thalli with highly variable abundances (0.35 to 65.6%, average of 9.0±16.5%, n=24); they were also detected in 21/24 supernatants but at 10-fold lower relative abundances (0.05 to 5.9%, average of 0.81±1.28%, n=24). This Gammaproteobacteria class was represented by 26 bacterial orders (20 <1%). Alteromonadales and Cellvibrionales were the orders most represented detected in 4/12 (0.95±2.15%) and 11/12 (3.7±2.9%) thalli of strain 010, and in 6/12 (11.3±23.3%) and 12/12 (1.6±2.9%) thalli of strain 06, respectively. Both Gamma-proteobacterial orders were also detected, although at lower abundances, in all corresponding supernatants (n=12, average value <0.9%) and all coral skeletons (n=8, 0.3±0.3% for Alteromonadales and 2.4±5.4% for Cellvibrionales). Acidimicrobiia (represented by the Microtrichales order) and Phycisphaerae (represented by the Phycisphaerales order) were detected in 22/24 and 18/24 *Ostreobium* thalli, with variable relative abundances of 0.8-25.6% and 0.03-20%, respectively. They were also detected, although at lower abundances and prevalences, in 19/24 and 13/24 supernatants (with relative abundances of 0.05-4.8% and 0.1-10.9%, respectively), and also within coral skeletons.

The Thermoleophilia class (represented by the Gaiellales order) was specific to thalli of strain 010 (clade P1), where it was detected in 11/12 samples (3.1±1.9%, n=12) compared to none in strain 06 (clade P14); it was also detected at trace level in strain 010 supernatants (5/12, 0.13±0.20%) and in coral skeletons (3/8, <0.5%).

The Deltaproteobacteria (former Polyangia, Myxococcia, Leptospirae and Desulfovibrionia classes, *cf* NCBI taxonomy browser) were represented by only the Nannocystales order in cultured thalli and supernatants of strain 010 at 40.2 psu (3/12 and 3/12 with relative abundances of 1.2±2.5% and 0.4±1.1%, respectively), and were undetected in strain 06. In contrast, higher diversity and abundances of Deltaproteobacteria were detected in coral skeletons, with representatives of four orders (Polyangiales, Myxococcales, Leptospirales and Desulfovibrionales) and higher cumulated relative abundances (6.8±12.7%, n=8).

Venn diagrams revealed the core bacterial fraction (core microbiota) of *Ostreobium* thalli, i.e. the ASVs shared by both genotypes and all three salinities (Figure 3B). The core bacterial fraction specific to each genotype (strain-specific core microbiota) and persistent across salinities was also visualized (Figure 3C and 3D). Bubble plots summarized the proportion of each core bacterial order or core ASV per category (Figure 6). The classification of shared ASVs, and their average relative abundances and prevalence among algal thalli, supernatants and environmental coral skeletons, are detailed in Supplementary Table 5.

**Figure 6:**
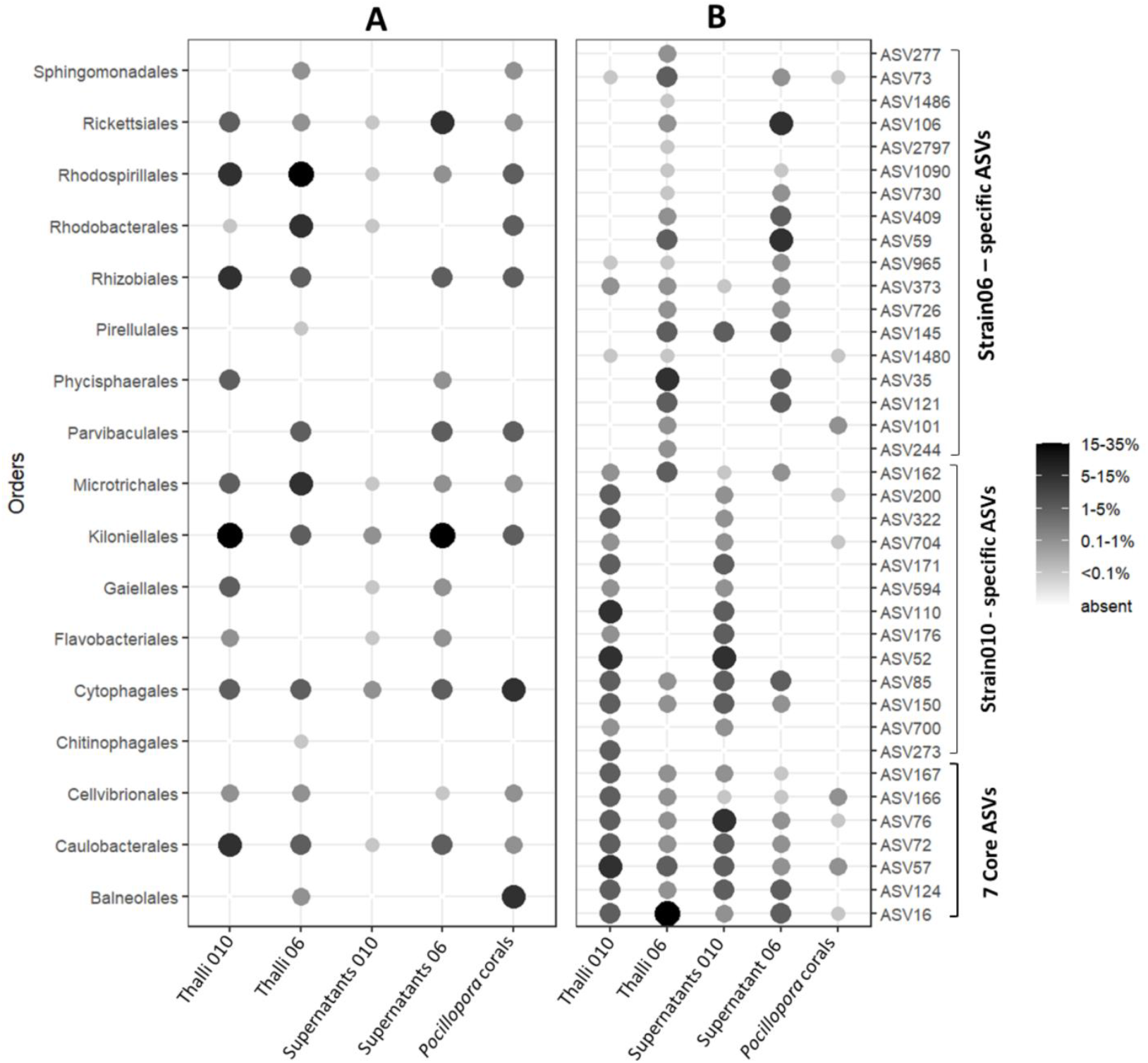
Core *Ostreobium* microbiota composition, shared at 3 salinities. The proportion of each (A) bacterial order or (B) ASV within a sample category is calculated as a percentage of total cumulated reads assigned to each category: thalli of 010 and 06, corresponding supernatants and environmental *Pocillopora* coral skeletons. Taxonomy classification after SILVA v138 SSU rRNA database released December 16, 2019. Abundances are grey-scale color and size coded.

Among the ASVs persistent across 3 salinities, only 7 bacterial ASVs were shared between thalli of both genotypes (1.8% out of a total of 400 bacterial ASVs for 010, corresponding to 18.6% cumulated abundance; 1.5% out of a total of 469 bacterial ASVs for 06, corresponding to 35.7% cumulated abundance; Figure 3B and Supplementary Table 5). This core microbiota was dominated by Alphaproteobacteria (5 ASVs), with one additional ASV belonging to Bacteroidia and another to Acidimicrobiia.

Within the Alphaproteobacteria, the numerically dominant and most prevalent taxon in *Ostreobium* core microbiota was ASV16 (unresolved at genus level, classified to the family Rhodospirillaceae, order Rhodospirillales), with 75-100% prevalence and relative abundances depending on algal genotype. It was 10 times more abundant in 06 (∼32 %) than in 010 (∼3%) (Figure 6 and Supplementary Table 5), confirming its PCA-detected contribution to strain differentiation (Supplementary Figure 4). Also present in culture supernatants, but at much lower relative abundances (<6.3% in 06 and <1% in 010) than in thalli, ASV16 was detected in only one coral skeleton, at trace level (<0.01% abundance). The second dominant alphaproteobacterial taxon was ASV57 classified to the family Hyphomonadaceae (order Caulobacterales), with 75-83% prevalence and 1.8 to 5.7% average relative abundances in 010 and 06, respectively (Figure 6 and Supplementary Table 5). It was detected at lower prevalence and relative abundances in supernatants and at trace levels and low prevalence in coral skeletons. Three additional core alphaproteobacterial taxa were detected at trace levels (<1%) in 06 and at 1.7-3.2% relative abundances in 010 (Figure 6 and Supplementary Table 5). The Rhizobiales order (reclassified as Hyphomicrobiales) was represented by ASV72 (classified to the genus *Labrenzia*), which was highly prevalent (91%) in both strains. It was also detected albeit at lower prevalence and abundances in supernatants and undetected in coral skeletons. The Kiloniellales order (reclassified as Kiloniellaceae family within the Rhodospirillales) was represented by ASV124 (affiliated to the genus *Fodinicurvata*), detected in both strains at all salinities and 33% - 66% prevalence. It was also present in supernatants (especially of strain 06) and undetected in coral skeletons. Finally, the Rickettsiales order was represented by ASV76 at low abundances (<2%) but high to medium prevalence in thalli of both strains (100% in 010 and 50% in 06). This ASV76 was classified to the basal lineage (AB1) of Rickettsiales defined via metagenome assembly of rare bacterial taxa in the marine bryozoan *Bugula neritina* (Miller *et al*., 2016). It was detected with same prevalence but higher abundances in culture supernatants and had low prevalence (25%) at trace level in *Pocillopora* coral skeletons.

Within the Bacteroidia class, the Cytophagales order was represented by ASV166 (classified to the genus *Candidatus Amoebophilus*, family Amoebophilaceae), which was the only core bacteria with 100% prevalence, being represented at low abundances (<2%) in all thalli of both strains. It was very rare in culture supernatants (2/24, with average abundances <0.02%), but persistently detected (75% prevalence) at trace levels in the skeletons of environmental *Pocillopora* corals. Finally, within the Acidimicrobiia class, the order Microtrichales was represented by the genus *Ilumatobacter*, with ASV167 persistent across salinities in both strains (50% prevalent in 06, 75% prevalent in 010) and abundances of 0.27 and 1.86% in 06 and 010, respectively.

Additionally, we detected strain-specific core microbiota (persistent across salinities, and additional to the 7 ASVs presented above). It was composed for strain 010 of 12 ASVs (3.0% out of a total of 400 bacterial ASVs; among which 4/12 ASVs were detected at trace levels <1%; Figure 3D and Supplementary Table 5) and for strain 06 of 19 ASVs (4.1 % out of a total of 469 bacterial ASVs; among which 13/19 ASVs were detected at trace levels <1%; Figure 3C and Supplementary Table 5). Several taxa were represented by multiple ASVs, with cumulated abundances depending on algal genotypes (Figure 6). Within Kiloniellales (reclassified to Kiloniellaceae), cumulated abundances of *Fodinicurvata* and *Pelagibius* were 18.9% in 010 *versus* only 1.3% in 06. Within Rhodospirillales, the Rhodospirillaceae family amounted to 5.8% in 010 *versus* 34.8% in 06. Within Rhizobiales, *Labrenzia*, unresolved Ahrensiaceae family members and *Devosia* were more abundant in 010 (8.3%) than in 06 (1.8%). Finally, within Microtrichales, cumulated abundances of *Illumatobacter* were 1.9% in 010 *versus* 5.5% in 06. Additional families had representatives only in the strain-specific core microbiota. Indeed, unresolved Rhodobacteraceae family members and *Paracoccus* were detected at 11.7% in 06 (<0.1% in 010). Distinct Cyclobacteriaceae representatives were detected in 010 (at 3.8% cumulated abundances) and in 06 (at 3.2% cumulated abundances). *Muricauda* Flavobacteriaceae were detected only in 010. The Phycisphaeraceae SM1A02 lineage and the Gaiellales (unresolved family members) were also detected only in 010 (∼3% each).

### *Genotype-driven structuration of* Ostreobium *microbiota, with limited salinity effect*

The classification of bacterial profiles by algal host genotype or abiotic, salinity factor was confirmed by sparse Partial Least Squares – Discriminant Analysis (sPLS-DA) of bacterial ASV relative abundances, and correlation analysis of sample projection coordinates with composition, to highlight informative ASVs that explain microbiota differentiation.

For the factor genotype, sPLS-DA results showed projection of bacterial communities into 2 clearly separated groups (Figure 7A, 6.57%, PC1), demonstrating that microbiota variability was structured by strain. All 4 most discriminant bacterial ASVs (4 selected ASVs with correlation coefficient >0.7 for projection on PC1) belonged to the core microbiota and were differentially expressed between 06 and 010. ASV16 (Rhodospirillaceae) was significantly more abundant in 06 than 010 (confirming results of PCA correlation circles plot, Supplementary Figure 4B). Alternately, 3 ASVs were more abundant in 010 than 06: ASV176 (Kiloniellaceae, confirming results of correlation circles plot of PCA at order level, Supplementary Figure 3), ASV171 (Cyclobacteriaceae), and ASV200 (Gaiellales) (Figure 7B).

**Figure 7:**
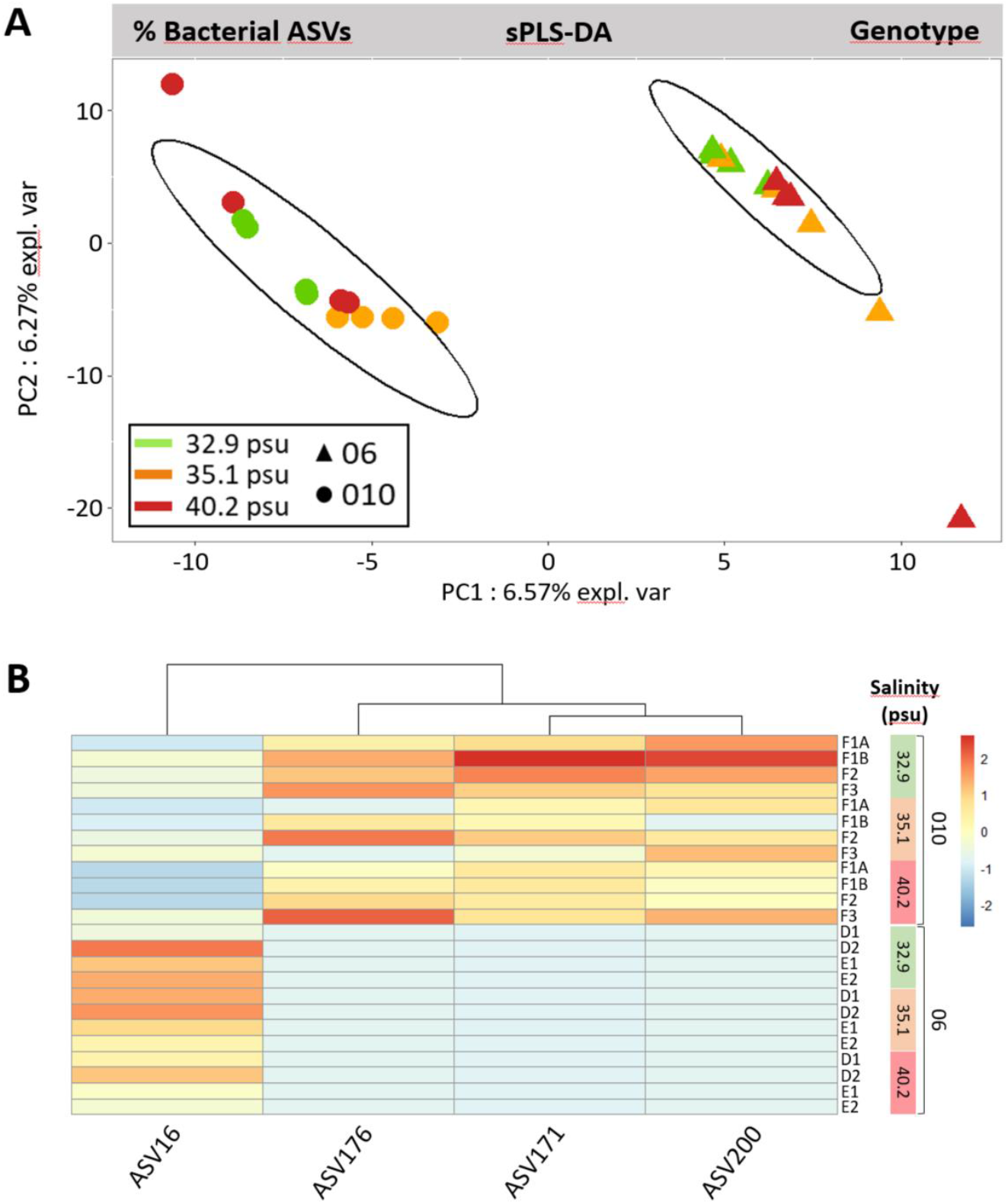
Algal genotype-driven structuration of bacterial ASVs communities in *Ostreobium* thalli. (A) sparse Partial-Least Square Discriminant Analysis (sPLS-DA with 95% confidence statistical ellipses). (B) Heatmap of bacterial ASVs with correlation coefficient >0.7 on Component 1 of the PLS model, with color-coded variation of relative abundances (red indicates more abundant bacterial ASV, salinity levels for each strain 010 or 06 are reported on the right).

For the factor salinity (analyzed separately for 06, Figure 8A; and 010, Figure 8B), sPLS-DA results showed that the microbiota of each strain (genotype) responded differently. For strain 06 (Figure 8A), there was a clear separation of the projections on PC1 (11.94% explained variability) of bacterial communities at high salinity (40.2 psu) *versus* intermediate (35.1 psu) and low (32.9 psu) salinities, which overlapped. For strain 010 (Figure 8B), it was the bacterial communities at low (32.9 psu) salinity that clearly projected on PC1 (13.47% explained variability), separately from intermediate (35.1 psu) and high (40.2 psu) salinities which overlapped. A single most discriminant ASV597 (Rhizobiales, affiliated to the genus *Devosia*), responsive to salinity increase, was shared by thalli of 06 and also 010 (more abundant at high salinity in both strains, Figure 8C). Additionally, in 010, four other ASVs were responsive to salinity increase (more abundant at high salinity), amongst which ASV57 (Caulobacterales Hyphomonadaceae from the core microbiota) in addition to ASV170, ASV383 and ASV976 (Figure 8C).

**Figure 8:**
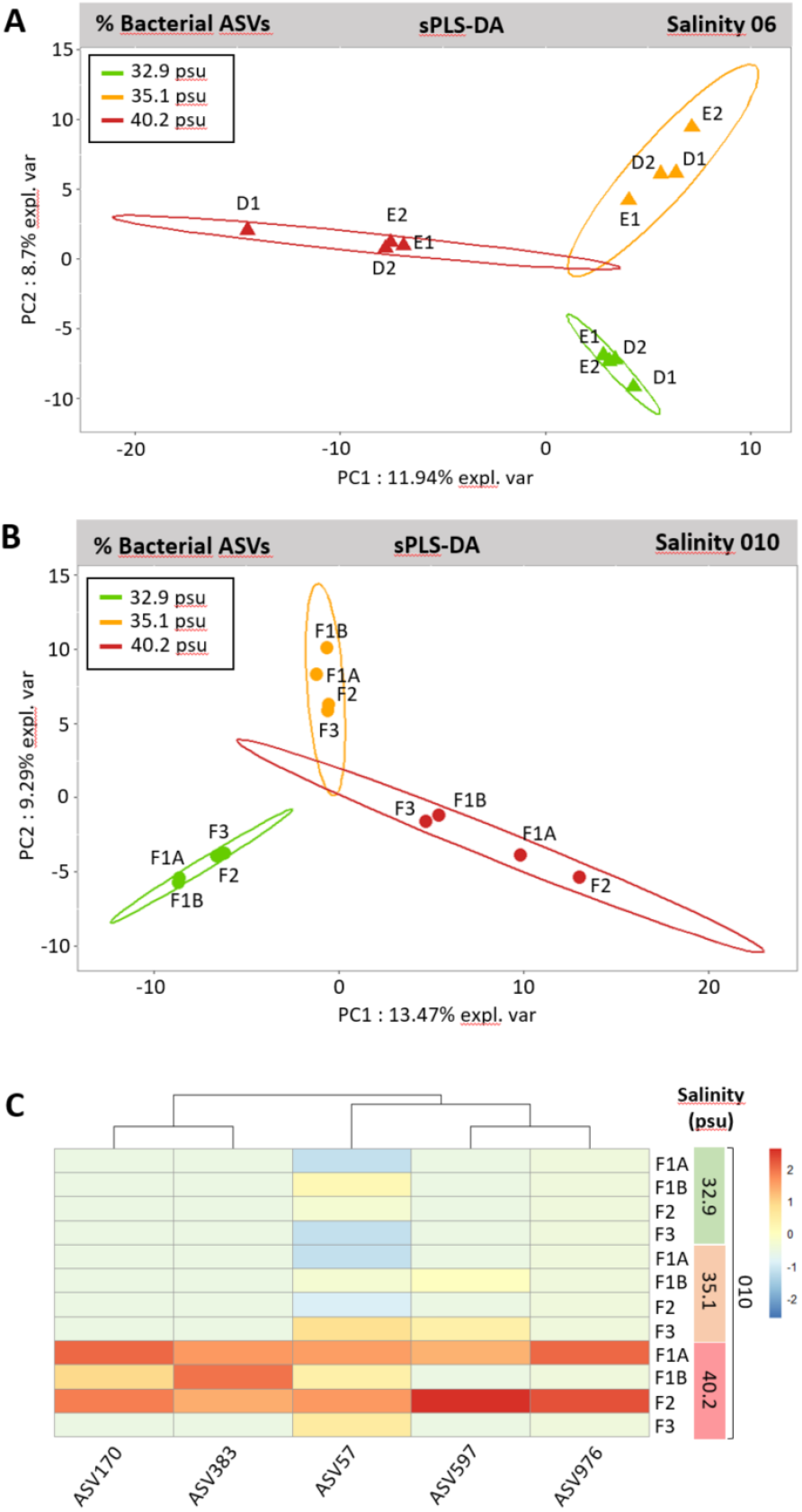
Secondary salinity structuration of bacterial ASVs communities in *Ostreobium* thalli, within each genotype. (A) Strain 06 (*rbc*L clade P14) and (B, C) strain 010 (*rbc*L clade P1). (A, B) sparse Partial-Least Square Discriminant Analyses (sPLS-DA with 95% confidence statistical ellipses, and replicate culture code names (FXX, EXX, DXX)). (C) Heatmap of the 010-associated bacterial ASVs with correlation coefficient >0.7 on Component 1 of the PLS model, with color-coded variation of relative abundances (red indicates more abundant bacterial ASV). For 06, only 1 bacterial ASV with correlation coefficient >0.7 on Component 1 of the PLS model was identified (ASV597, enriched at 40.2 psu).

## Discussion

*Ostreobium* is a cryptic but ubiquitous Ulvophyceae partner resident in the coral holobiont, with a yet unexplored phycosphere. This study is the first to characterize *Ostreobium* bacterial microbiota composition and to address its variability at different salinities. For this, we used a reductionist approach based on cultured strains to pull apart the microbial complexity of *Ostreobium*-colonized microniche coral skeleton environments. The bacteria co-cultured with strains of 2 distinct *Ostreobium* genotypes co-isolated from an aquarium-grown *Pocillopora* sp. coral colony (Massé *et al*., 2020) displayed partial overlap with *in hospite* endolithic bacteria of environmental *Pocillopora* corals from Guam and the Red Sea. Comparison of multiple cultures revealed a small-size core microbiota, shaped by algal genotype and with limited response to salinity increase.

### *Bacterial diversity reduction in domesticated* versus *wild* Ostreobium *(within corals)*

Overall, a reduction in Amplicon Sequence Variant (16S rDNA ASV) richness was observed in “domesticated” *Ostreobium* unialgal strains maintained in culture collections, compared to “wild” *Ostreobium*-colonized *Pocillopora* coral endolithic assemblages from Guam and Eilat. In coral-associated endolithic bacterial assemblages, systemic surveys highlight the co-occurrence of prokaryotes with several *Ostreobium* genotypes together with fungal and protists microeukaryotes (Sweet *et al*., 2011; Marcelino *et al*., 2017a; Cárdenas *et al*., 2022). The endolithic bacterial community is thus shaped both by taxonomic composition of this large diversity of residents and by environmental variables, which fluctuate in time and between colonies and reef sites. This complexity does not allow to determine which bacterial fraction is specifically associated with which endolithic algae. Our reductionist experimental approach provides simplified *in vitro* inoculum (unialgal *Ostreobium* cultures) and long-term controlled environmental variables (here, salinity) that revealed bacterial communities consistently associated to specific *Ostreobium* genotypes. A similar approach has been used for Symbiodiniaceae microalgal endosymbionts of coral tissues, to identify their core bacteria in multiple cultures of long-term propagated strains (Lawson *et al*., 2018) and in response to temperature stress (Camp *et al*. 2020).

Here, we present new data on skeleton bacteria from Red Sea (Eilat) and central Pacific (Guam) *Pocillopora* sp. corals, completing previous surveys of *Pocillopora damicornis* from Papua New Guinea (Marcelino *et al*., 2017b) and from the Great Barrier Reef (Ricci *et al*., 2022). Our results confirm the presence of the genus *Endozoicomonas* (Gammaproteobacteria; 14 ASVs detected, 1.2% cumulated abundances), which is a prevalent symbiont in many corals (Neave et *al*., 2017). *Endozoicomonas* is especially abundant and persistent within tissues of Red Sea *Pocillopora verrucosa* (2 phylotypes, 90% cumulated abundance; Pogoreutz *et al*., 2018). It was detected but at much lower abundances in the skeleton of *Pocillopora damicornis* (Marcelino *et al*., 2017b; Ricci *et al*., 2022) and 9 other scleractinian species (where it represented 7% of skeleton ASVs *versus* 33% of tissue ASVs across the whole data set; Ricci *et al*., 2022).

Our results highlight partial overlap between *Pocillopora* coral endolithic bacteria and bacteria co-cultured with *Ostreobium* strains, initially isolated from *Pocillopora acuta*. Domesticated *Ostreobium* strains thus contained a fraction of co-isolated endolithic bacteria, able to survive in long-term aerobic cultures. This observation validates the use of *Ostreobium* strains as a simplified model for *in vitro* studies of the phycosphere of endolithic algae. Deltaproteobacteria were among the difficult bacterial taxa to maintain *in vitro*, represented by only one sparse order (Nannocystales) in thalli of *Ostreobium* strain 010 cultured at high salinity, in contrast to four abundant orders in *Pocillopora* skeletons. Of note, the genus *Endozoicomonas* detected in coral skeletons was not detected in *Ostreobium* cultures of either strain, suggesting absence of association with *Ostreobium* algae in our culture conditions.

The bacterial communities were highly variable across multiple cultures of the same *Ostreobium* strain. This finding is consistent with the high variability reported in cultures of the green microalga *Tetraselmis suecica* strain F&M-M33 (Piampiano *et al*., 2019), where opportunistic bacteria may be enriched or introduced and carried over during successive subcultures separated from initial inoculum. Here, the initial inoculum for *Ostreobium* cultures was a tuft of interwoven filaments, with oxygen and nutrient gradient likely developing from periphery to center, driving spatial microbiota differentiation. Subsampling by random initial fragmentation is likely to account for the variability detected here in subsequent replicate *Ostreobium* cultures.

### *Small size core* Ostreobium *bacterial microbiota, structured by algal genotype*

Both algal genotypes shared core bacterial taxa, found consistently across all three salinities, corresponding to 7 ASVs classified to 7 distinct families, with cumulative abundances ∼19% for 010 and ∼36% for 06. This small subset of ASVs (1.5-1.8% of the detected ASVs in each strain) is in agreement with studies of another siphonous Bryopsidale, *Caulerpa* sp., in which the core microbiota across four host species was <1% of the total detected taxa (41 OTUs shared across 4 *Caulerpa* species; Aires *et al*., 2015).

*Ostreobium* associated bacteria were dominated either by Kiloniellales or Rhodospirillales depending on host genotype. A tight association (epilithic/endophytic) can be suggested for Rhodospirillales ASV16 (unresolved Rhodospirillaceae), Caulobacterales ASV57 (unresolved Hyphomonadaceae), Rhizobiales ASV72 (*Labrenzia*), Cytophagales ASV166 (*Candidatus ‘Amoebophilus’*) and Microtrichales ASV167 (*Illumatobacter*), based on differential abundance patterns, 3 to 10 fold relatively more abundant in thalli than corresponding supernatants. For these core taxa, limited detection may be carried-over in supernatants via DNA amplification from potential residual *Ostreobium* fragments retained on filters.

The 16S rDNA sequence of Rhodospirillaceae ASV16 (unclassified at genus level), was the most abundant core microbiota member, and matched with that of an uncultured Rhodospirillaceae bacterium clone from the Caribbean *Orbicella (Montastrea) faveolata* coral (100% identity with JQ516442.1; Kimes *et al*., 2013). Multiple additional Rhodospirillaceae were also detected in 06 (44 ASVs) and 010 (39 ASVs). This finding is congruent with previous detection of dominant sequences attributed to 2 Rhodospirillaceae species (∼55-60% relative abundances) in the microbiome of the siphonous green alga *Caulerpa ashmeadii* in the Gulf of Mexico (Sauvage *et al*., 2019).

Four other core alphaproteobacterial taxa shared a functional category related to nitrogen metabolism, based on function prediction hypotheses that can be inferred from the metabolism of cultured type strains of Fodinicurvataceae (Kiloniellales), Hyphomonadaceae (Caulobacterales), and Rhizobiales. *Ostreobium* typically colonizes nitrogen-limited environments, such as biomineralized CaCO3 skeletons of corals (with <1% dry weight of organic matter), and its epilithic filaments and free-living propagules are also adapted to N-limited oligotrophic reef seawater. Bacteria-assisted nitrogen metabolism may support algal holobiont metabolism, and its adaptation to N-depleted environments.

Kiloniellales (re-classified as Kiloniellaceae) have been involved in overall N metabolism in the context of their association with *Caulerpa prolifera*, another Bryopsidale alga (Aires *et al*., 2015). Here, *Ostreobium* thalli (especially of strain 010) contained abundant Kiloniellales, especially at high salinity. Fodinicurvataceae are halophilic bacteria, abundant in intertidal sandflat algal mat environments (Huang *et al*., 2020), with type strains of the *Fodinicurvata* genus isolated from salt mine sediments (Wang *et al*., 2009). The nitrate-reducing strain *Fodinicurvata halophila*, isolated from a marine saltern, contains C18:1ω7c as a major fatty acid (36%; Infante-Dominguez *et al*., 2015), which was also detected in our *Ostreobium* strains (Massé *et al*., 2020). Kiloniellaceae have recently been documented in the context of the coral holobiont, within skeleton assemblages of several coral species (Ricci *et al*., 2022). However, the nature of association of Fodinicurvataceae with *Ostreobium* still needs to be investigated, due to the high sequence abundances noted here in the supernatants of cultured strains, suggesting lifestyle traits of opportunistic bacteria amplified by the *in vitro* conditions (nutrient-enriched culture medium).

Nitrate-reducing, strictly aerobic bacteria in the family Hyphomonadaceae (here represented by ASV57 in the order Caulobacterales) are adapted to oligotrophic environments in the ocean (Abraham and Rohde, 2014) and were already detected in marine macroalgae, including in the kelp *Nereocystis luetkeana* (Weigel and Pfister, 2021), the seagrass *Halophila stipulacea* (Weidner *et al*., 2000), and the red alga *Porphyra yezoensis* (Fukui *et al*., 2013).

Within Hyphomonadaceae, the species *Algimonas porphyrae* isolated from red alga *Porphyra yezoensis* is characterized by high content of the fatty acid C18:1ω7c (Fukui *et al*., 2013). As noted above, this monounsaturated fatty acid has previously been reported to be relatively abundant in coral-isolated *Ostreobium* strains (Massé *et al*., 2020), providing biochemical evidence supporting the occurrence of Hyphomonadaceae as *Ostreobium* core bacteria.

Finally, the genus *Labrenzia* (order Rhizobiales) was identified here as a core *Ostreobium* bacteria, in agreement with previous occurrence reports in 5 *Bryopsis* species (Bryopsidales), kept in long-term unialgal laboratory cultures (Hollants *et al*., 2011) or sampled across Atlantic, Mediterranean and Pacific localities, at salinities ranging from 32.4 to 37.6 psu (Hollants *et al*., 2013). *Labrenzia* bacteria have also been detected in the core microbiota of cultured Symbiodiniaceae dinoflagellates, which are major endosymbionts of coral tissues (Lawson *et al*., 2018). Members of Rhizobiales are known to fix dinitrogen (N2) into dissolved ammonium and nitrate, transferred to their plant host, promoting its growth (Carvalho *et al*., 2014). We propose that nitrogen-fixing Rhizobiales bacteria could play an important role in the nutrition and growth of *Ostreobium* strains. Benthic turf algae form a reef compartment that includes carbonate-colonizing *Ostreobium* algae, and are known to fix nitrogen (Roth *et al*., 2020). Overall, the functional diversity of *Ostreobium* associated bacteria should be further studied for potential significance in reef nitrogen and carbon cycling.

Another important member of *Ostreobium* core bacteria was *Ilumatobacter* (order Microtrichales). This genus was first described from bacteria isolated from beach sand off the Sea of Japan, and is also associated with marine sponges (Funjinami *et al*., 2013). Here, its persistent detection in thalli of *Ostreobium* strains 06 (3 ASVs) and 010 (2 ASV), and minor detection in supernatants, suggests tight association. The genome of the sponge-isolated cultured strain contains biosynthetic enzymes for bacterial vitamins (B6, K), omega-3 fatty acids and xanthin carotenoids, and lacks several aminoacid biosynthetic enzymes, but encodes adhesion microbial surface components (Fujinami *et al*., 2013). Similar putative pathways might be conserved in *Ostreobium-*associated *Ilumatobacter*, that might serve to adhere to algal host filaments.

Additionnally, within the core bacteria specific to strain 010, the Phycisphaerales ASV322 SM1A02 lineage was detected. Phycisphaerales are known for their ability to break down and metabolize complex algal polysaccharides (Boedeker *et al*., 2017) that might be produced specifically by the 010 lineage. Phycisphaerales were also detected in the culture supernatants of strain 010, supporting the hypothesis of bacterial utilization of potential algal-secreted polysaccharidic cell wall and mucilage components.

Altogether, the bacterial assemblages revealed here can be considered as extended phenotypes of each *Ostreobium* genetic lineage, supporting distinct metabolism, such as the distinct N and C assimilation patterns and fatty acids metabolism previously recorded in 010 and 06 (Massé *et al*., 2020). Core bacterial associates may be selected by their capacity to use specific algal metabolites and secreted mucilage. Reciprocally, associated bacteria may contribute limiting N, nutrients and vitamins for algal growth.

### *Potential intracellular symbionts of* Ostreobium

Intracellular bacteria are a still underexplored fraction of the coral holobiont and its associated Symbiodiniaceae and Ostreobineae microscopic algae (reviewed by Maire *et al*, 2021b). Here, we document for the first time within the core *Ostreobium* bacteria, 16S rDNA sequences of 2 potential intracellular symbionts.

*Candidatus ‘Amoebophilus’* ASV166 (family Amoebophilaceae; order Cytophagales; class Bacteroidia), was found to be ubiquitously associated with all algal thalli, 75% of *Ostreobium*-colonized *Pocillopora* coral endolithic assemblages and absent from almost all supernatants. The *Candidatus ‘Amoebophilus’* lineage includes obligate intracellular bacteria with reduced genome, discovered within a marine amoeba (Schmitz-Esser *et al*., 2010).

Recorded as a rare bacterial member of the coral holobiont (Ainsworth *et al*., 2015) it was recently found at high abundances in the skeleton microbiome of several coral species (e.g. *Isopora prolifera* Ricci *et al*., 2021, *Pocillopora damicornis* and Poritidae; Ricci *et al*., 2022). Its persistent detection in all 12 samples of both *Ostreobium* genotypes, in all 3 salinities, reveals a persistent association and suggests close and possibly intracellular association with *Ostreobium* filaments.

The Rickettsiales_AB1 lineage (ASV76) was also persistently detected in the thalli of both *Ostreobium* strains. Rickettsiales are obligate intracellular bacteria with reduced genomes, known to infect animals but also algae. Indeed, intracellular Rickettsiales were sequenced and visualized in the unicellular green alga *Carteria* (Volvocales) (Kawafune *et al*., 2012) and they have been detected in 2 *Bryopsis* sp. species (Bryopsidales), sampled in the Pacific and North Sea at 18.0 and 32.4 psu (Hollants *et al*., 2013). Intracellular Rickettsiales were also detected in coral host tissue (*Candidatus ‘*Aquarickettsia rohweri’; Klinges *et al*., 2019), and in two Symbiodinaceae strains, *Cladocopium goreaui* and *Durusdinium glynii*, isolated from the corals *Acropora tenuis* and *Porites lobata*, respectively (Maire *et al*., 2021a). The Rickettsiales_AB1 lineage highlighted here within cultured *Ostreobium* thalli was also detected in a similar study of multiple Symbiodiniaceae cultures, propagated from a monoalgal isolate from *Acropora tenuis* coral (Buerger *et al*., 2022). Here, ASV76 was equally or more abundant in supernatants than in *Ostreobium* thalli, and undetected in *Pocillopora* endolithic assemblages. Its presence in supernatants in addition to thalli might be explained by parasitism on the dead portion of filaments (without Dapi-stained nuclei) and/or utilization of N nutrients provided by decaying cells and culture medium.

### *Limited* Ostreobium *bacterial community adjustments to salinity increase*

Differentiation in microbiota profiles was detected between low (32.9 psu) and high (40.2 psu) salinities in both algal genotypes, with an intercalation of the intermediate salinity profile either with low (for strain 06) or high (for strain 010) salinity. Each strain (genotype) responded differently, with a seemingly higher salinity threshold for bacterial community changes in 06 than 010, and a common proportion of increased Rhizobiales ASV597. Overall, the *Ostreobium* microbiota salinity response was highly variable, showing a flexibility that may support salt tolerance of this alga. Indeed, in other models, the composition and function of algal-associated bacterial microbiota has an essential role in the health of their host and its adaptation to changes in salinity (Singh and Reddy, 2016). In the brown alga *Ectocarpus*, microbiota shifts observed in response to lower salinity were shown to be necessary for algal host survival (Dittami *et al*., 2016). In the cultured Ulvophyceae model *Ulva mutabilis*, the algal low salinity response involved interactions with marine bacteria *Roseovarius* and *Maribacter* (Ghaderiardakani *et al*., 2020). The flexibility of the bacterial microbiota, shown here in long-term acclimatized algal cultures, may be part of the mechanisms that facilitate adaptation of *Ostreobium* to a large salinity range, supporting its ubiquitous distribution in carbonate reefs. This hypothesis supports a recently proposed concept of the contribution of microbiome flexibility to accelerate the acclimatization and adaptation of holobionts to environmental change (Voolstra and Ziegler, 2020).

### *Epiphytic* versus *endophytic bacterial associations with* Ostreobium *filaments*

Here, *in situ* hybridization (CARD-FISH) targeting 16S rRNA with universal bacterial probes allowed the first visualization of metabolically active bacteria in or near *Ostreobium* filaments (in the extracellular mucilage). Our study builds on a previous FISH visualization of bacteria in the apical fragments of a related freshwater *Bryopsis* alga (Hollants *et al*., 2011). However, we were unable to morphologically differentiate spatial microniches at the algal thallus level in these free-living *Ostreobium* growth forms (outside the carbonate habitat). Indeed, the orientation relative to a carbonate bioerosion front was absent in the thallus, composed of a tuft of branched interwoven filaments (Massé *et al*., 2020, Pasella *et al*., 2022). The siphonous cytology of microscopic *Ostreobium* thalli, with cytoplasmic streaming in septae-lacking tubes, presents a challenge for subsampling of individual filaments, for culture propagation or regionalized microbiota characterization. Thallus fragmentation involves high risk of spilling the contents of broken filaments into surrounding culture medium, which is a source of artifactual “supernatant” bacteria. Orientation should be optimized, so that bacteria can be precisely localized to the filament structures. For example, it has been shown in macroscopically well differentiated Caulerpa (Bryopsidale) thallus (fronds/rhizoids) from the Mediterranean Sea that distinct epiphytic and endophytic bacterial communities existed (Aires *et al*., 2013 and 2015), spatially differentiated from the rhizobiome (Morrissey *et al*., 2019). Here, at the spatial resolution of our epifluorescence study, we could not unambiguously differentiate epiphytes from endophytes among the bacterial phylotypes.

Future CARD-FISH investigations will use taxon-specific probes designed from the sequences obtained in this study, to visualize their distribution within *Ostreobium* thalli. Targeted taxa will include abundant Rhodospirillaceae and Kiloniellaceae, and rare but widespread bacteria with putative intracellular lifestyles, such as *Candidatus Amoebophilus* and the Rickettsiales_AB1 lineage.

### Extension to other Ostreobium strains and other bacterial markers

To further investigate the core *Ostreobium* bacteria, additional studies are needed on other algal strains, for example isolated from Great Barrier Reef corals (Pasella *et al*., 2022), to verify microbiota similarities between different host lineages spanning the whole phylogenetic diversity of the Ostreobineae suborder (4 *tuf*A-based clades *sensu* Marcelino and Verbruggen, 2016, or families *sensu* Sauvage *et al*., 2016). Future research should also involve bioeroding growth forms, complementary to the free-living growth forms studied here, to investigate potentially different selection of bacterial associates in the more nutrient-limited and less oxygenated carbonate habitat, supporting *Ostreobium*’s changing metabolism across habitats (Massé *et al*., 2020).

Finally, bacterial community diversity characterization is well known to be biased by selected DNA extraction method, molecular markers, and amplification primers. Here, the selected bacterial 16S rRNA gene region V5-V7 limited co-amplification of algal plastidic 16S rDNA as in brown macroalgae (Vieira *et al*., 2016; Tourneroche *et al*., 2020) and *Caulerpa* (Morrissey *et al*., 2019). Yet, amplification of the *tufA* elongation factor which is used for metabarcoding of *Ostreobium* algae (Marcelino and Verbruggen, 2016) also yielded non-target bacterial *tuf*A sequences that were not detected via conventional 16S rDNA metabarcoding. Amplification of *tuf*A sequences from heterotrophic bacteria in addition to phototroph cyanobacteria and *Ostreobium* algae was previously reported (Sauvage *et al*., 2016). Our preliminary results with *tuf*A detected rare representatives of the Nitrospira phylum that remained undetected via V5-V7 16S rDNA metabarcoding (1 ASV classified to Nitrospiraceae detected per *Ostreobium* strain, in biomass extracts from 5 out of 7 strains - 06, 017, 018B, 018C, and 019, not detected in 010 and 018A, described in Massé *et al*., 2020).

Another rare alphaproteobacterial taxon was revealed by *tuf*A sequencing: ASVs classified to the Xanthobacteraceae family were persistently detected at low abundance in 6/8 strains (1 ASV in 06, 1 ASV in 010, 2 ASVs in 017, 2 to 4 ASVs in 018A, and 1 ASV in 018B). Hence, the *Ostreobium*-associated bacterial diversity retrieved here cannot be considered exhaustive, and is likely to be further extended by the future use of complementary markers.

**In conclusion**, this study of *Ostreobium* bacterial taxonomic diversity under experimental salinity stress showed that the microbiota of long-term-propagated strains was structured primarily by algal genotype, with limited influence of salinity between 32.9 and 40.2 psu. The identification of 7 core bacterial taxa persisting across salinities and genotypes paves the way for further investigations of their role in the physiology and overall health of their algal host. More generally, the plasticity of microbiota composition highlighted here likely contributes to *Ostreobium*’s high ecological tolerance, supporting its ubiquitous distribution and resilient colonization of carbonates in changing reef salinities, and advocates for more in-depth approaches to their roles and importance to the coral holobiont.

## Acknowledgments

The authors thank Dr. Marc Gèze and Mr. Cyril Willig of the Centre de microscopie de Fluorescence et d’IMagerie numérique (CeMIM) of the Muséum national d’Histoire naturelle (MNHN) for advice in the acquisition of fluorescence images. Students Joëlle Robbe and Maëva Goulais helped extract coral skeletons DNA and amplify *tuf*a and *rbc*L sequences. The authors also thank Mrs. Amandine Blin from the MNHN-CNRS UAR2700 ‘Acquisition et Analyse de Données pour l’Histoire naturelle’ (Pole Analyse de Données) for advice on data visualization and Pr. Alain Paris from the MNHN-CNRS UMR7245 for his help on statistical analyses.

## Authors Contributions

A.M. conceived the research, acquired funding, and designed the experimental protocols with the help of I.D-C. and S.D. J.D. performed the experiments. I.D-C. helped for CARD-FISH experiments and acquired fluorescence microscopy image. A.M., J.D. analyzed 16S rDNA sequence dataset. A.C.W. and A.M. analyzed *tuf*A and *rbc*L sequence dataset. S.D. and C.D. helped analyze and discuss the 16S rDNA metabarcoding data. A.M. and I.D-C. wrote and reviewed the manuscript, which was commented by all authors.

