## Supplementary Figure 1 for "Bacterial microbiota of *Ostreobium*, the coral-isolated chlorophyte ectosymbiont, at contrasted salinities"

**Supplementary Figure 1:** Experimental design for multiple cultures and salinity treatments of two *Ostreobium* strains. m: month(s).

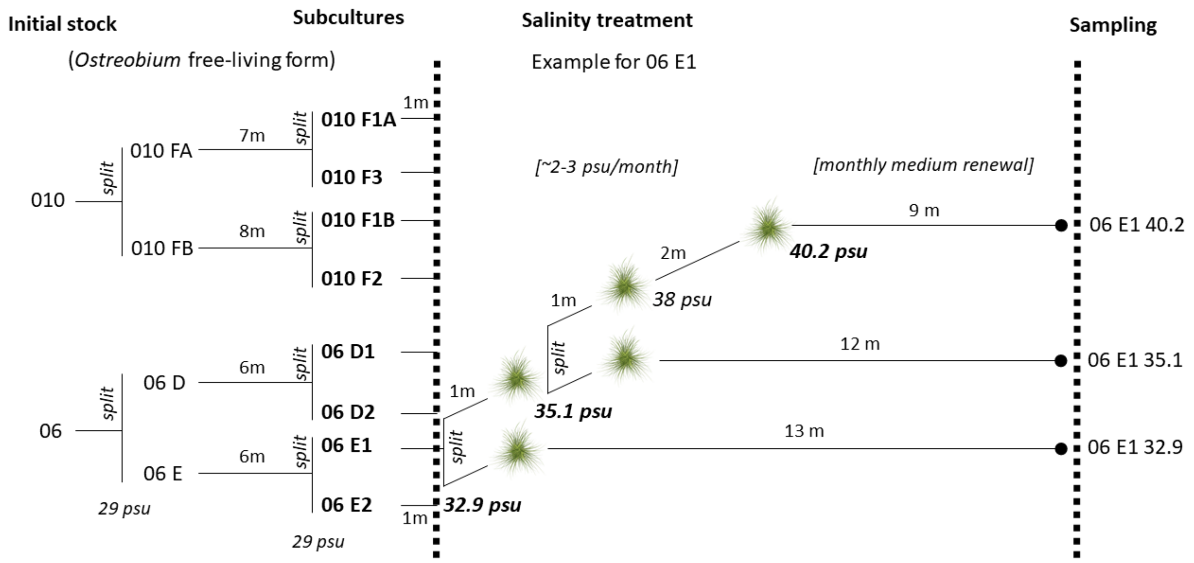
