## Supplementary Figure 2 for "Bacterial microbiota of *Ostreobium*, the coral-isolated chlorophyte ectosymbiont, at contrasted salinities"

**Supplementary Figure 2:** Rarefaction curves from V5-V7 region of bacterial 16S rDNA amplified from cultured *Ostreobium* thalli and supernatants, and environmental *Pocillopora* coral skeletons. Curves were constructed using the rarefied dataset, after filtering out contamination (internal) controls.

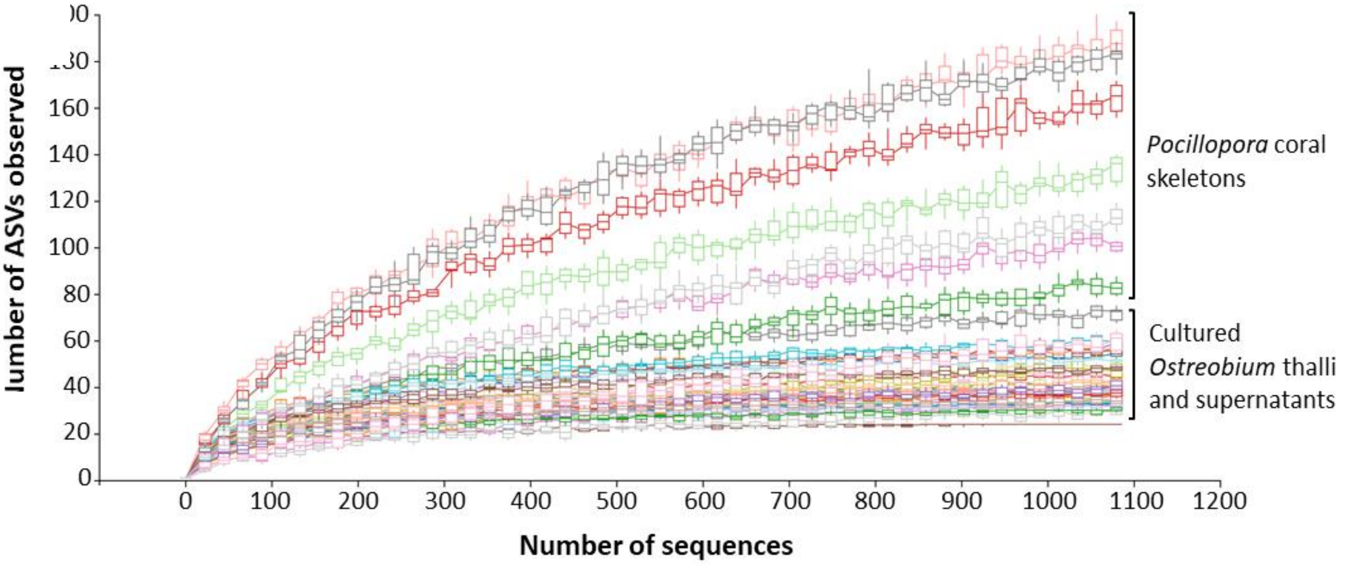
