## Supplementary Figure 3 for "Bacterial microbiota of *Ostreobium*, the coral-isolated chlorophyte ectosymbiont, at contrasted salinities"

**Supplementary Figure 3:** Structure of the bacterial communities visualized by Principal Component Analysis at order level (unrarefied and untransformed proportions data). (A) Partial overlap between communities of cultured *Ostreobium* thalli, their supernatants and those of coral skeletons. (B) Clustering of *Ostreobium* thalli bacterial community profiles by algal genetic lineage (010: *rbc*L clade P1, 06: *rbc*L clade P14) more than salinity. (B1) Score plot (with 2 outlier cultures 06_E2_35.1 psu and 06_E2_40.2 psu) with indication of culture code names (FXX, EXX, DXX); (B2) loading plot of PCA analysis.

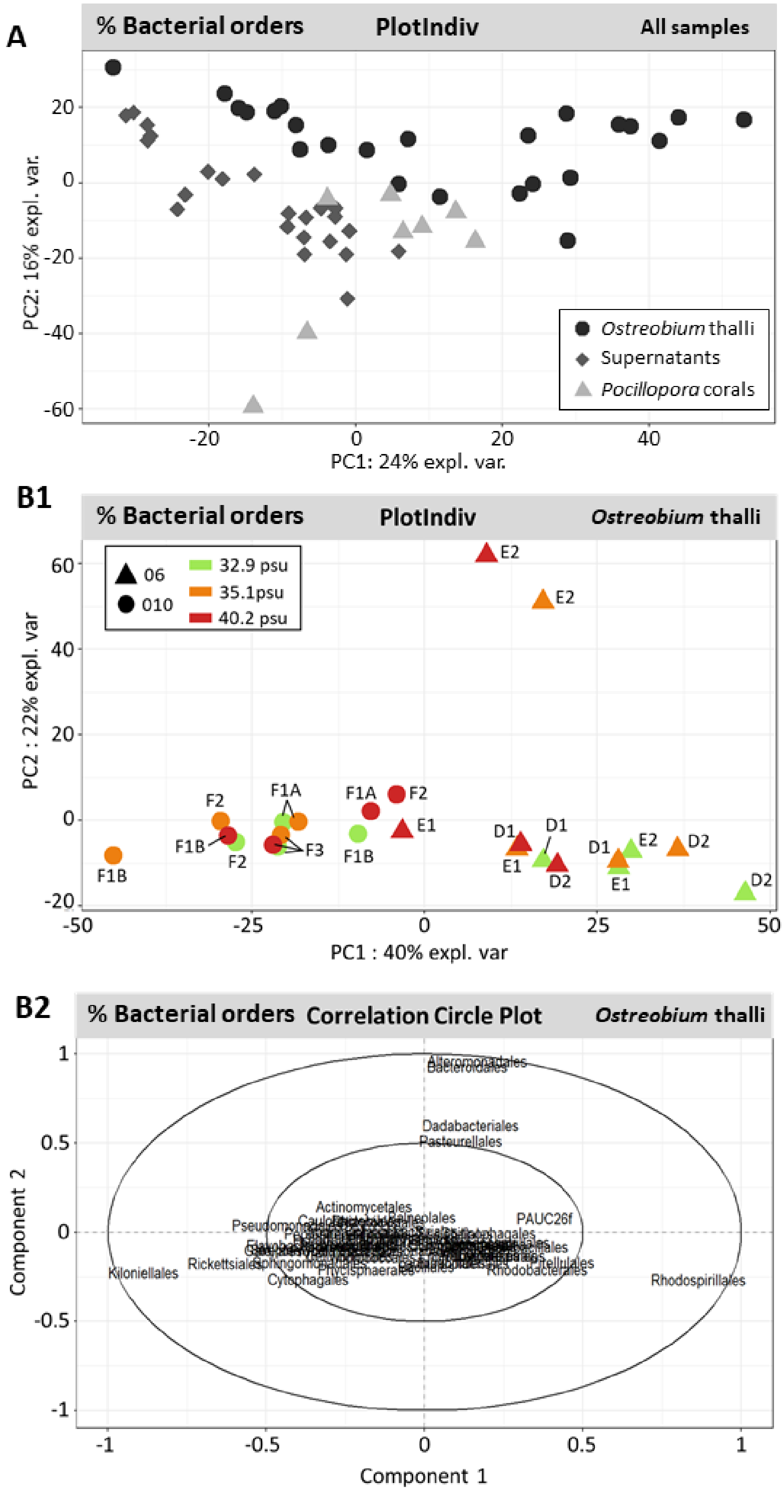
