## Supplementary Figure 4 for "Bacterial microbiota of *Ostreobium*, the coral-isolated chlorophyte ectosymbiont, at contrasted salinities"

**Supplementary Figure 4**: Correlation circle plot of PCA analysis (at ASV level; illustrated in Figure 4) for (A) all samples (cultured *Ostreobium* thalli, supernatants and environmental coral skeletons) and (B) *Ostreobium* thalli of each genotype.

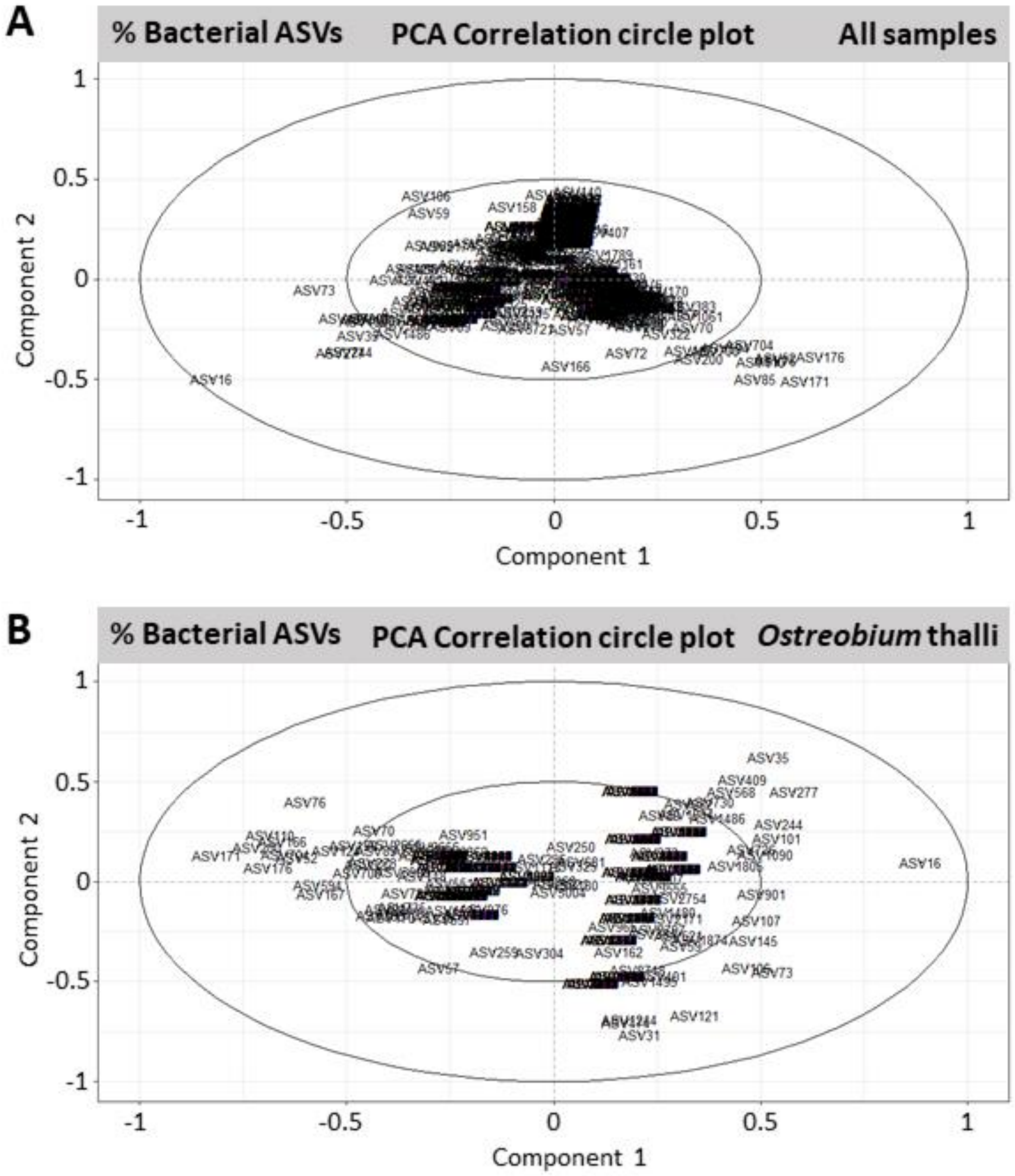
