## Supplementary Figure 5 for "Bacterial microbiota of *Ostreobium*, the coral-isolated chlorophyte ectosymbiont, at contrasted salinities"

**Supplementary Figure 5**: Taxonomic composition of bacterial classes in cultured *Ostreobium* thalli and supernatants, and the environmental *Pocillopora* coral skeletons. Salinity and algal genotype or reef site are indicated below each individual sample column. The classes detected at trace levels were aggregated in the <1% abundance category.

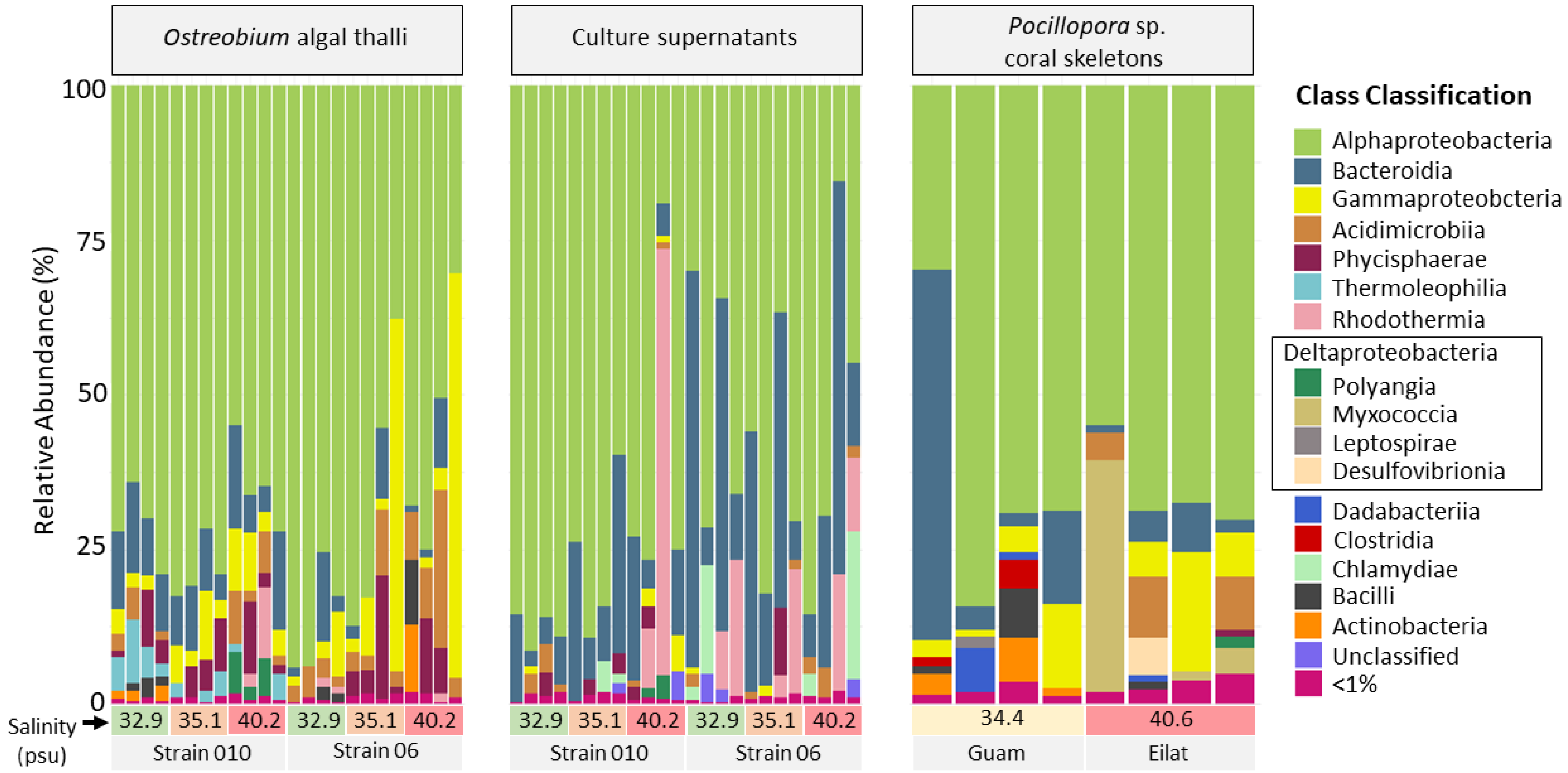
