## Supplementary Table 1 for "Bacterial microbiota of *Ostreobium*, the coral-isolated chlorophyte ectosymbiont, at contrasted salinities"

**Supplementary Table 1**: (A) Read counts of 16S rDNA ASVs obtained from cultured *Ostreobium* thalli, their supernatants, coral skeletons and internal controls using Illumina MiSeq sequencing (unfiltered data). (B) Read counts of bacterial 16S rDNA ASVs after removing ASVs affiliated to Eukaryota, Chloroplast, Mitochondria, Unassigned ASVs and all ASVs of the internal controls (filtered data). (C) Fasta sequences of 16S rDNA ASVs (unfiltered data). (D) Taxonomy of bacterial 16S rDNA ASVs (filtered data). (E) Fasta sequences of Ostreobineae *tuf*A ASVs detected in coral skeletons from Guam (Pacific) and Eilat (Red Sea).

This Supplementary Table 1 is available on figshare with doi: 10.6084/m9.figshare.21953075
