## Supplementary Table 4 for "Bacterial microbiota of *Ostreobium*, the coral-isolated chlorophyte ectosymbiont, at contrasted salinities"

**Supplementary Table 4:** Summary of pairwise permutational multivariate analysis of variance ('adonis' function) of Bray-Curtis distances (n=999 permutations, $\alpha = 0.05$) for pairs of factor levels. Bray-Curtis distances were calculated from arcsine square root transformed unrarefied bacterial ASV proportions. (A) Variance comparison between environmental *Pocillopora* coral skeletons *versus in vitro* algal cultures (*Ostreobium* thalli and supernatants). (B) Variance comparison between cultured algal categories (*Ostreobium* thalli *versus* supernatants), genetic lineages (010 *rbc*L clade P1 *versus* 06 *rbc*L clade P14), salinities (32.9 *versus* 35.1 *versus* 40.2 psu) and their interactions. (C) Pairwise comparisons of variance within algal cultures for the salinity factor (p-adjusted with Bonferroni correction).

**A) Variance comparison between environmental coral skeletons and *in vitro* algal cultures**

| | Df | SumOfSqs | R2 | F | Pr(>F) |
|---|---|---|---|---|---|
| **metadata$treatment** | 1 | 1.2663 | 0.0533 | 3.0400 | 0.001* |
| **Residual** | 54 | 22.4939 | 0.9467 | NA | NA |
| **Total** | 55 | 23.7603 | 1 | NA | NA |

**B) Variance comparison *in vitro* algal culutres between sample nature (thalli *vs* supernatants), genotype (010: *rbc*L clade P1 vs 06: *rbc*L clade P14), salinity (32.9, 35.1, 40.2 psu) and their interactions**

| | Df | SumOfSqs | R2 | F | Pr(>F) |
|---|---|---|---|---|---|
| **metadata$nature** | 1 | 0.3553 | 0.0185 | 1.0121 | 0.425 |
| **metadata$genotype** | 1 | 1.3594 | 0.0708 | 3.8729 | 0.001* |
| **metadata$salinity** | 1 | 1.9946 | 0.1039 | 5.6825 | 0.001* |
| **metadata$salinity:metadata$nature** | 1 | 0.2319 | 0.01208 | 0.6607 | 0.949 |
| **metadata$salinity:metadata$genotype** | 1 | 0.8469 | 0.0441 | 2.4128 | 0.003* |
| **metadata$genotype:metadata_2$nature** | 1 | 0.2056 | 0.0107 | 0.5857 | 0.976 |
| **metadata$salinity:metadata$genotype:metadata$nature** | 1 | 0.1573 | 0.0082 | 0.4482 | 0.998 |
| **Residual** | 40 | 14.0402 | 0.7316 | NA | NA |
| **Total** | 47 | 19.1911 | 1 | NA | NA |

**C) Pairwise comparison of salinity factor in *in vitro* algal cultures**

| | pairs | Df | SumsOfSqs | F.Model | R2 | p.value | p.adjusted |
|---|---|---|---|---|---|---|---|
| 1 | 32.9 *vs* 35.1 | 1 | 1.0077 | 2.7601 | 0.0843 | 0.001* | 0.003 |
| 2 | 32.9 *vs* 40.2 | 1 | 2.0866 | 6.0346 | 0.1675 | 0.001* | 0.003 |
| 3 | 35.1 *vs* 40.2 | 1 | 0.9783 | 2.5245 | 0.0776 | 0.001* | 0.003 |
